# BioVault: A privacy-first data visitation platform for equitable global collaboration in biomedicine

**DOI:** 10.64898/2026.02.12.705603

**Authors:** Manaswitha Edupalli, Tivadar Péter Török, Madhava Jay, Keelan Jordan, Zeynab Abdirahman, Devy Frederick, Carika Weldon, Rana Dajani, Xi Dawn Chen

## Abstract

Biomedical datasets representing diverse populations are essential for advancing precision medicine, yet remain siloed due to regulatory, sovereignty, and privacy constraints. Existing data-sharing solutions remain limited. Centralized repositories and Trusted Research Environments (TREs) require data migration into external infrastructure, introducing governance complexities, cost barriers, and constraints on accountable reuse. Secure computation frameworks from modern cryptography, and federated learning models demand deep domain expertise and substantial engineering effort, limiting their accessibility in resource-constrained environments. These barriers disproportionately impact under-resourced institutions, limiting equitable participation in global collaborations. Here, we introduce BioVault (https://www.biovault.net), an open-source platform for privacy-first biomedical collaboration through a peer-to-peer data visitation network, where analysis code travels to data rather than data being transferred to centralized systems. BioVault can be deployed as a desktop application or command-line interface, enabling out-of-the-box use for participants in diverse resource settings. BioVault supports both clinical and research workflows, and with built-in permissioning, audit trails, and local governance controls, it enables data holders and collaborators to retain oversight and control while participating in collaborative research. We demonstrate BioVault’s utility in enabling two cross-border collaborations with under-resourced communities: a genome-wide association study of Type 2 Diabetes in Circassian and Chechen populations in Jordan, and allele frequency estimation across Caribbean cohorts. In both cases, analyses were executed locally without data export. We further demonstrate compatibility for user-defined secure, federated computation protocols built on state-of-the-art cryptographic tools (multiparty computation and homomorphic encryption), by lowering barriers to their deployment. Together, these results establish BioVault as a general-purpose framework for decentralized analytics that reduces the technical and financial constraints to biomedical collaboration – enabling diverse institutions to participate as equal partners in global discovery, while preserving privacy and data sovereignty.

## Introduction

Biomedical research increasingly depends on the responsible sharing of large-scale biomedical datasets to advance precision medicine^1–3^. Cross-institutional aggregation of genomic, clinical, and molecular data enables the discovery of novel disease variants^4,5^, improves prediction performance of machine learning models^6^, and reveals disease mechanisms across populations^7^. However, the expansion of data generation has coincided with increasingly stringent data protection laws, data sovereignty requirements, and privacy safeguards that restrict cross-border transfer of human data. This growing misalignment has created a structural bottleneck: scientific progress requires global data collaboration, yet regulatory and governance constraints limit such collaboration and slow the translation of biomedical data into clinical benefit^8^.

Existing data-sharing architectures remain limited. Centralized repositories such as dbGaP and the European Genome-phenome Archive (EGA) provide controlled access to archived datasets but require transfer of sensitive data to external infrastructure, introducing jurisdictional complexity, operational cost, and concentration of security risk^9–11^. Trusted Research Environments (TREs) enable remote analysis and reduce uncontrolled redistribution, but similarly depend on data migration into hosted platforms that are often controlled by well-resourced institutions or commercial vendors and weaken durable linkage between data owners and downstream analytical artifacts^12^. Federated data platforms seek to preserve local data control, but often require prior data harmonization and pose high infrastructural costs^13–17^. Secure computation frameworks enable privacy-preserving computation while maintaining local data control, but require deep expertise in cryptography, distributed algorithms, and domain-specific knowledge of biomedical analyses to implement successfully^18–21^. There is also a non-trivial engineering burden that is required to deploy such systems in real-world environments^22,23^. In practice, enabling collaborative research frequently entails either data movement across jurisdictions, centralized custodianship, or significant technical overhead, reinforcing a perceived trade-off between privacy, sovereignty, and scientific participation.

These architectural constraints disproportionately impact emerging biobanks, small clinical centers, and under-resourced research groups– especially those in low- and middle-income countries (LMICs) and Indigenous communities– which face persistent barriers to participating as partners in global biomedical discovery^24–26^. The burden is most acute in resource-limited settings, where infrastructure and funding for sophisticated platforms are unavailable. Consequently, datasets collected from underrepresented populations are often exported for off-site analysis, and once copied into external infrastructure, institutions may lose practical control over how data are accessed, reused, and governed, limiting local capacity building and equitable benefit sharing from biomedical discovery^27–30^.

Here we introduce BioVault (https://www.biovault.net), an open-source platform for decentralized, privacy-first biomedical collaboration using data visitation via a peer-to-peer network (**Fig. 1A, B**). In BioVault, analytical code and model parameters travel to locally-governed datasets, and raw data never leaves institutional boundaries; only approved results are returned after local execution. To lower barriers to adoption, BioVault is designed for ease of use and is available as both a desktop application and a command-line interface, for out-of-the-box deployment. BioVault supports diverse biomedical data modalities and analytical workflows, including single-cell analysis, machine learning model training on physiological data, radiology imaging model inference, and clinical genomics analyses. We demonstrate BioVault’s utility in fostering global partnerships in two real-world settings: a genome-wide association study of Type 2 diabetes in Circassian and Chechen populations in Jordan and allele frequency analysis across Caribbean populations. In both cases, analyses were configured and executed locally by non-technical individuals without data export, using the user-friendly desktop interface. We further demonstrate compatibility with secure multiparty computation frameworks via Sequre^31^/Shechi^32^, enabling participants to deploy custom privacy-preserving workflows within BioVault’s architecture. Together, these results establish BioVault as a general-purpose, accessible platform for data visitation and federated analytics that can lower barriers to equitable collaboration, while maintaing privacy and data sovereignty.

**Figure 1:**
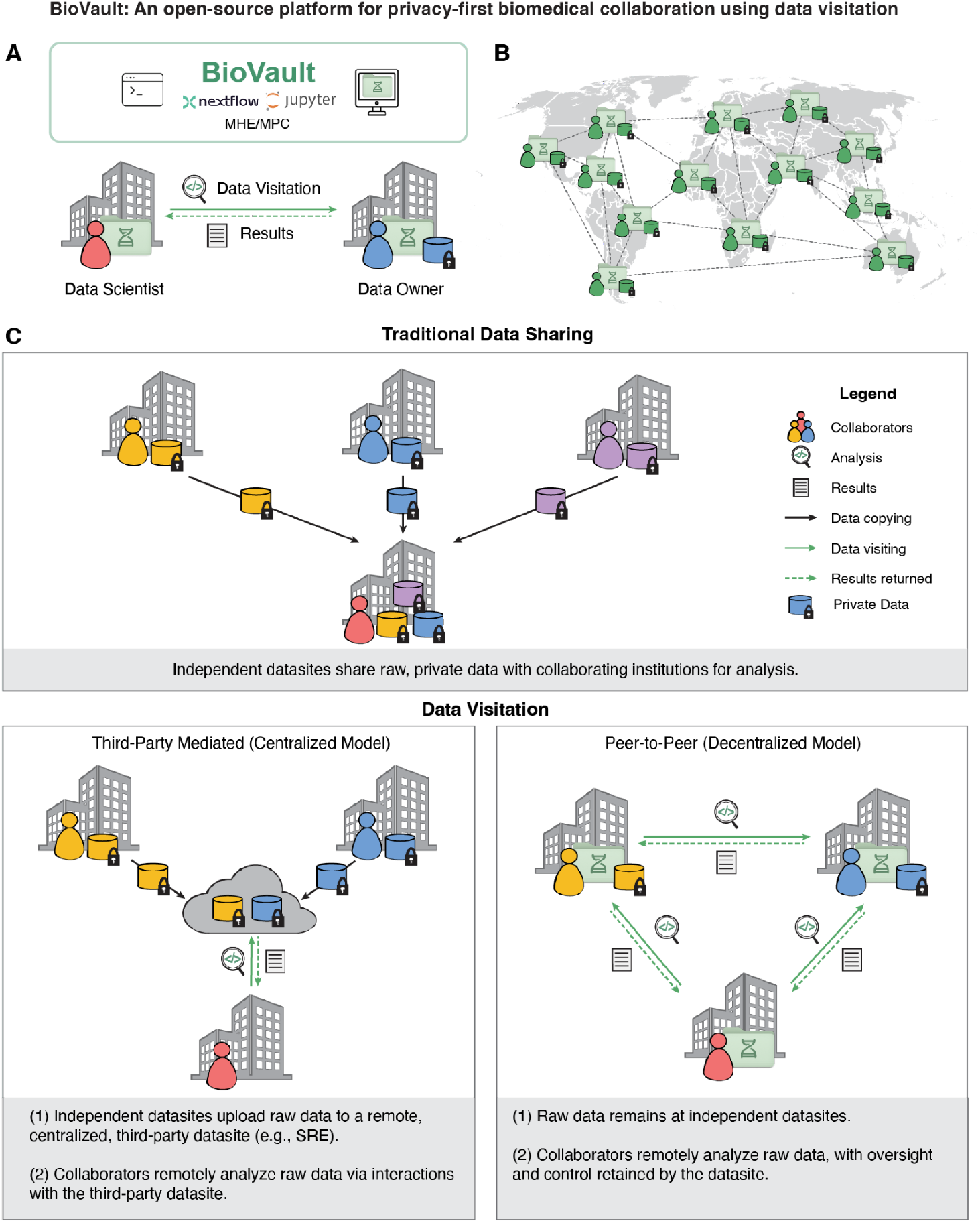
BioVault enables privacy-first biomedical collaboration through data visitation via a peer-to-peer network. **(A)** Overview of BioVault’s data visitation workflow. Under the peer-to-peer data visitation model, data scientists submit analysis code to data owners rather than accessing raw data directly. Data owners publish synthetic mock datasets that mirror the structure of private data otherwise not accessible, enabling data scientists to develop and validate pipelines before execution on real data (Supplementary Fig. 2). Raw data never leaves the data owner’s environment; only results from approved executions are returned. **(B)** BioVault operates across a decentralized network of independently governed datasites spanning institutions and jurisdictions, enabling global collaboration without centralized data pooling or cross-border data transfer. **(C)** Comparison of data sharing paradigms. In traditional data sharing (top), independent datasites share raw, private data directly with collaborating institutions. In third-party mediated data visitation (bottom left), raw data is uploaded to a centralized datasite, typically a secure research environment, where collaborators can analyze data remotely, but data owners have limited oversight over analyses performed. In the peer-to-peer data visitation model, proposed by BioVault (bottom right), raw data remains at independent datasites – collaborators can still analyze remotely, but with oversight and control retained by the data owner.

## Results

### BioVault enables privacy-first biomedical collaboration using peer-to-peer data visitation

BioVault provides a lightweight, decentralized infrastructure for privacy-first biomedical collaboration using data visitation, where analysis code travels to data rather than data moving into centralized systems (**Fig. 1A**). Available as both a desktop application (compatible with Linux, MacOS, and Windows) and a command-line tool (CLI), BioVault supports analysis pipelines written as Nextflow workflows or Jupyter Notebooks (**Supplementary Fig. 1**). To enable integration with emerging AI-driven research workflows, both the CLI and desktop application expose agent-accessible interfaces.

BioVault introduces a peer-to-peer network of independently governed environments for ease of privacy-first coordination and collaboration across borders and jurisdictions (**Fig. 1B**). It is built on SyftBox^33^, an open source protocol for secure, decentralized coordination of remote computation across independently governed environments (**Supplementary Note 1**). This technology allows BioVault to offer authenticated communication, execution request management, and result return between data scientists and data owners. Syftbox integration (Methods) also allows BioVault to treat linked data as generic files or databases under local control^34^, independent of data modality or analytical task, enabling researchers to configure the platform once by linking local datasets for reuse across heterogeneous workflows. Data is kept at its source, and messages and results are end-to-end encrypted before being sent. By providing a simple file-backed interface, we provide maximum compatibility and flexibility with existing data sources and future-proof the system for emerging AI capabilities.

To enable pipeline development against secure datasets that cannot be directly accessed, BioVault adopts a “twin” paradigm in which data owners publish a synthetic or mock dataset that mirrors the structure or schema of their private data without revealing sensitive content (**Supplementary Fig. 2, Supplementary Note 2**). These mock datasets allow researchers to develop and validate analysis pipelines prior to execution. Validated pipelines are then submitted as execution requests over the SyftBox network, where data owners review and authorize queries and execution in accordance with local policies. All operations are logged with transparent audit trails, and data owners retain granular control over permitted analyses and shared outputs (Methods). Computation is performed locally on the private datasets within the data owners’ environments. Only explicitly authorized results are returned. This approach supports deployment across heterogeneous settings, including personal laptops, institutional servers, high-performance computing systems, and cloud resources, without requiring data centralization. The resulting human-in-the-loop but AI-capable, attribution-based execution model aligns with existing access control, audit, and governance frameworks, enabling institutions to attribute computations, regulate outputs, and selectively participate in collaborations while preserving data sovereignty.

This peer-to-peer architecture distinguishes BioVault from existing biomedical data-sharing paradigms (**Fig. 1C**). In traditional data sharing, raw data is copied across institutional (and potentially jurisdictional) boundaries for analysis. Third-party-mediated models, such as Trusted Research Environments (TREs), reduce uncontrolled redistribution by centralizing data within managed cloud infrastructure, but may limit data owners’ oversight and create vendor dependence. In some instances, TREs sever the direct relationship between data owners and collaborators, potentially constraining the implementation of more advanced end-to-end privacy-enhancing technologies (PETs). In contrast, BioVault’s peer-to-peer data visitation model retains raw data at independent datasites while enabling remote analysis, with oversight and control maintained by the data owner (**Fig. 1C**, bottom right).

Together, these design choices establish BioVault as a flexible, decentralized platform for data visitation-based biomedical computation that lowers technical and operational barriers to participation while preserving local control over sensitive data. By enabling analyses to travel to data rather than requiring transfer into centralized environments, BioVault can support scalable collaboration across heterogeneous regulatory, institutional, and resource settings, while remaining agnostic to data modality, scientific domain, and analysis workflow.

### BioVault supports diverse clinical and biomedical workflows across data modalities

To demonstrate BioVault’s flexibility, we applied it to a set of representative use cases spanning diverse biomedical analyses and machine learning workflows involving sensitive clinical and patient-derived data. These use cases included patient-derived single-cell RNA-seq (scRNA-seq) data, physiological time-series data, large-scale clinical medical imaging, and personal genomic data from a rare disease patient. Across all workflows, data scientists submitted analyses, model training, or inference requests that were executed remotely within data owners’ environments. Raw data was never transferred to data scientists; only explicitly approved outputs were returned. While we are focused on biomedical applications, it is clear that these same paradigms are relevant to many fields in science that would benefit from access to private datasets.

#### Remote single-cell analysis on patient-derived molecular data

We first leveraged BioVault to enable data visitation of patient-derived scRNA-seq data from a metastatic breast cancer cohort^35^ (30 patients, >157k cells) (**Fig. 2A**). scRNA-seq data are biologically rich and typically require iterative, exploratory analysis to interpret. At the same time, they are highly sensitive: single-cell count matrices can enable linking attacks that leak identifiable genetic and phenotypic information of individuals, even when shared in summarized or pseudobulk form^36^. In this use case, the data scientist developed and iteratively refined an analysis workflow using synthetic mock datasets in an interactive Jupyter environment, while the data owner retained custody of the private data and approved execution requests. Approved commands were executed locally within the data owner’s environment, including standard preprocessing steps, quality controls, and downstream outputs (e.g., UMAP embeddings), without exposing raw single-cell gene expression profiles. By structuring the workflow as a sequence of approved execution steps with returned intermediate outputs using standard Python tooling, both parties can iteratively refine the analysis while maintaining local governance and control. This demonstrates that role-separated data visitation via BioVault supports complex, interactive, exploratory single-cell analyses while minimizing privacy risk.

**Figure 2:**
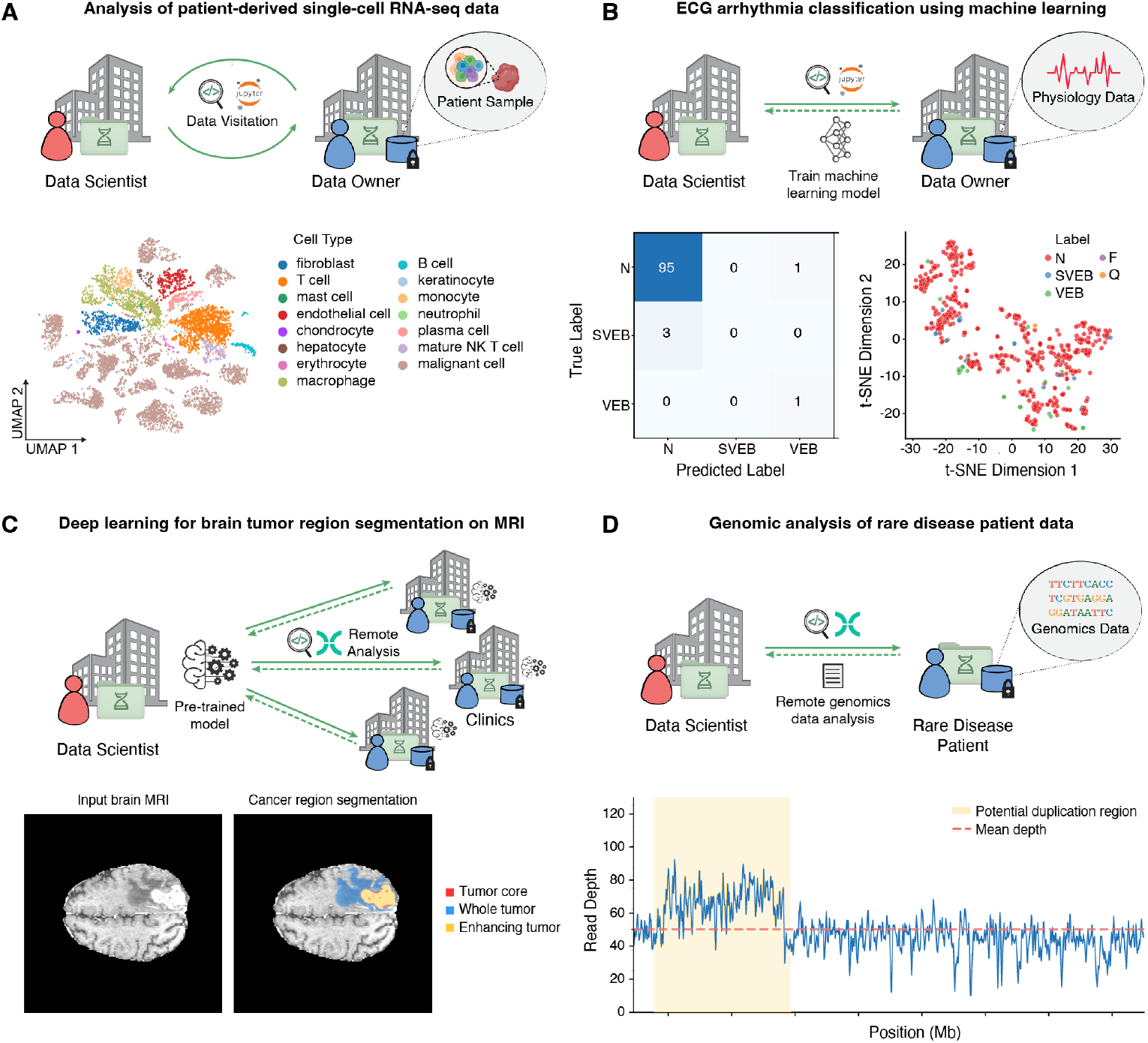
BioVault supports remote analysis of diverse biomedical data types. **(A)** BioVault enables remote preprocessing and exploratory analysis of patient-derived single-cell RNA-seq under data visitation, matching standard single-cell workflows while preserving privacy. Shown is a UMAP embedding of metastatic breast cancer biopsy cells annotated by cell type. **(B)** Remote training of an ECG arrhythmia classifier on the PhysioNet MIT-BIH dataset. Training code is developed and validated on mock data, then executed by the data owner to fit the model without exposing underlying ECG signals. Shown are the confusion matrix and a t-distributed stochastic neighbor embedding (t-SNE) of learned representations for normal (N), fusion (F), supraventricular ectopic beat (SVEB), ventricular ectopic beat (VEB), and unknown (Q) **(C)** Model-to-data inference for brain tumor region segmentation on MRI using a pretrained network. Example input slice and predicted segmentation are shown, with regions labeled as whole tumor (blue), enhancing tumor (yellow), and tumor core (red). **(D)** Remote analysis of rare disease whole-genome sequencing returns a locus-restricted read-depth profile for CNV interpretation, highlighting a potential duplication (shaded). Across all analysis, training, or inference requests are executed within the data owner’s environment. Raw data stays local and private, and only explicitly approved outputs are returned.

#### Remote training of machine learning models on physiological time-series data

Next, we showcase BioVault’s ability to support remote machine learning model training without data centralization using physiological time-series data from the MIT-BIH Arrhythmia dataset obtained from PhysioNet ^37,38^ (**Fig. 2B**). In this demonstration, the dataset was hosted within an emulated data-owning environment implemented as a virtual machine. Relevant features were previously extracted from electrocardiogram (ECG) signals to classify regular and irregular heartbeats. Model architectures and training code were developed and validated using synthetic data, then executed remotely within the data-owning environment. Only trained model parameters and performance metrics were returned to the data scientist, without exposing underlying ECG signals. To help data scientists evaluate how model architectures and hyperparameters perform on real data, live logs and training metrics can be provided by the data owner’s side during training sessions. This workflow demonstrates that BioVault enables compute-intensive, privacy-first model development on sensitive physiological data and could support the deployment of diagnostic models across distributed hospital networks while maintaining institutional control over patient data.

#### Deep learning-based model-to-data inference for brain tumor region segmentation on large-scale MRI data

Machine learning models are increasingly being used to analyze medical images in clinical and research settings, but the size and sensitivity of imaging data often limit their transfer outside of hospital systems. We utilized BioVault to perform remote deep learning inference for brain tumor region segmentation from the Multimodal Brain Tumor Image Segmentation Benchmark^39^ (BraTS) on large-scale, multimodal, magnetic resonance imaging (MRI) data (**Fig. 2C**). Transfer of this data is constrained by technical and governance considerations common to clinical environments, necessitating the need for data visitation. A pretrained neural network was shared with the data-owning environment to perform inference on MRI scans. Raw imaging data remained local, while only derived segmentation outputs and quantitative features were returned to the data scientist. This use case demonstrates that BioVault enables model-to-data inference on large, sensitive medical imaging datasets without requiring data movement, consistent with established clinical data governance practices.

#### Privacy-first analysis of personal genome data for rare disease investigation

Finally, we employed BioVault on whole-genome sequencing data from an already diagnosed rare disease patient carrying a clinically relevant copy number variant (CNV), analyzed under explicit participant consent (**Fig. 2D**). Raw sequencing data was locally held within the data-owning environment, while alignment and downstream analyses were executed remotely under a data visitation framework. Analysis pipelines were developed and validated using mock data and then approved for execution on the patient’s genome, producing only a region-restricted pileup and read-depth profile spanning the CNV locus of interest. These derived outputs enabled validation and interpretation of the copy number alteration without exposing the full genome or unrelated variants. This example demonstrates that BioVault supports clinically meaningful rare disease analyses on high-fidelity raw data while strictly limiting data egress to narrowly scoped, interpretation-ready results.

Together, these use cases demonstrate that BioVault is a flexible, general-purpose platform for data visitation that supports a wide range of biomedical analyses, including exploratory data analysis, remote machine learning model training, and model-to-data inference. Across molecular, physiological, imaging, and genomic domains, BioVault enables computation on sensitive data without requiring data movement, while preserving analytical fidelity and local control. These results establish BioVault as a domain-, data-, and analysis-agnostic framework for privacy-first biomedical collaboration.

### BioVault enables GWAS of Type II Diabetes in Jordan via data visitation

To evaluate BioVault’s capacity to enable cross-border genomic analyses while preserving data sovereignty, we established a collaboration between datasites in Jordan and the United States (**Fig. 3A**). Historically, similar studies have required the transfer of biological samples or individual-level genomic data to external institutions for sequencing and analysis, raising concerns regarding data sovereignty and limiting local analytical capacity. Using BioVault, however, we performed a remote genome-wide association study (GWAS) of Type II Diabetes (T2D) in Circassian and Chechen populations from Jordan through data visitation, providing an alternative to centralized data-sharing models.

**Figure 3:**
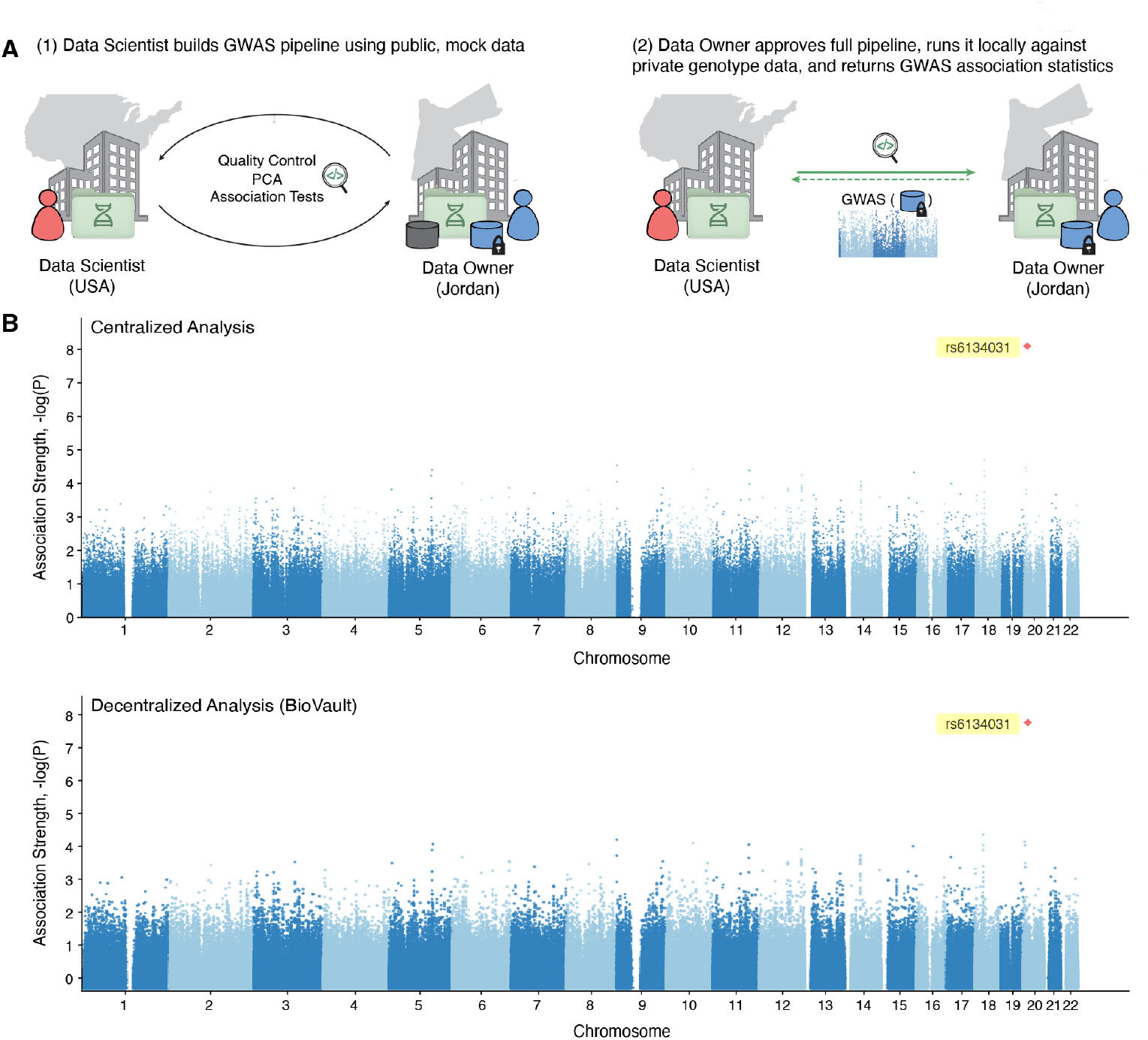
Cross-border GWAS on Jordanian genotype data using data visitation. **(A)** Schematic of the BioVault-enabled cross-border GWAS workflow for type II diabetes (T2D) in Chechen and Circassian populations from Jordan. Analysis steps for each phase of GWAS: from applying quality control filters, to computing principal components to accounting for population stratification, and association statistic testing. Each phase was developed and validated based on synthetic mock genotype data. Each GWAS step was then submitted as an execution request to the data owner to be run locally on private data. Only final association statistics were returned; no individual-level data (including genotypes, phenotypes, or covariates) were transferred outside the data-owning institution. **(B)** Manhattan plots comparing case-control GWAS results for T2D on Circassian and Chechen populations (n = 284; 67 cases, 217 controls; 627,744 autosomal SNPs) obtained from centralized analysis with direct data access (top) and BioVault-enabled remote analysis (bottom). Logistic regression was performed with the top 10 principal components as covariates to account for population stratification. The *P*-values were calculated using a two-sided score-based test and were not adjusted for multiple testing. Results replicate previously identified T2D-associated loci in these populations, demonstrating concordance between remote and centralized approaches while preserving privacy. All analysis steps were carried out using PLINK v1.9, either remotely via BioVault or locally in the centralized approach.

Previous GWAS of T2D have been conducted predominantly in populations of European and East Asian ancestry^40–42^. The Circassian and Chechen populations, with their distinct genetic background and endogamous structure, offer a valuable opportunity to characterize population-specific genetic risk factors for T2D in an underrepresented group.

Each stage of the GWAS pipeline was developed and validated using synthetic mock genotype data that mirrored the structure of the private dataset, including quality control, principal component analysis, and association testing (**Fig. 3A**). Once validated, the complete GWAS pipeline was submitted as an execution request and run remotely within the local computing environment of the datasite in Jordan. Throughout the analysis, genotyping data remained under local institutional control, and only final association test statistics were returned. No individual-level data, including genotypes, phenotypes, and covariates such as sex and age, were transferred outside of Jordan.

Post quality-control, the final dataset consisted of approximately 280 Circassian and Chechen individuals and 630K SNPs. Principal component analysis was performed to capture population structure, and the top 10 principal components were included as covariates in logistic regression-based association testing using PLINK v1.9 (Methods). GWAS results from the BioVault-enabled remote analysis closely match those from a centralized analysis that had sustained access to private, individual-level data throughout pipeline development (**Fig. 3B**).

Our findings replicated previously identified T2D-associated SNP locus (rs6134031) in the Circassian and Chechen populations^43^ (**Fig. 3B**). Together, these results demonstrate BioVault’s utility in enabling genomic discovery while maintaining data protection (privacy) and data sovereignty.

### BioVault enables cross-border analysis of allele frequencies across Caribbean populations

To evaluate BioVault’s ability in enabling cross-border population genetic analyses via data visitation while preserving data sovereignty, we initiated a remote allele frequency analysis from the United States, with computations executed remotely by collaborators in Saint Lucia and Bermuda (**Fig. 4A, Supplementary Note 3**). The Bermuda data site hosted genotype data from multiple Caribbean cohorts, including the Bahamas, Barbados, the British Virgin Islands, and Trinidad and Tobago. Per-country SNP-array callsets were analyzed locally, and only aggregate allele frequency statistics were returned, without transfer of individual-level genotype data.

**Figure 4:**
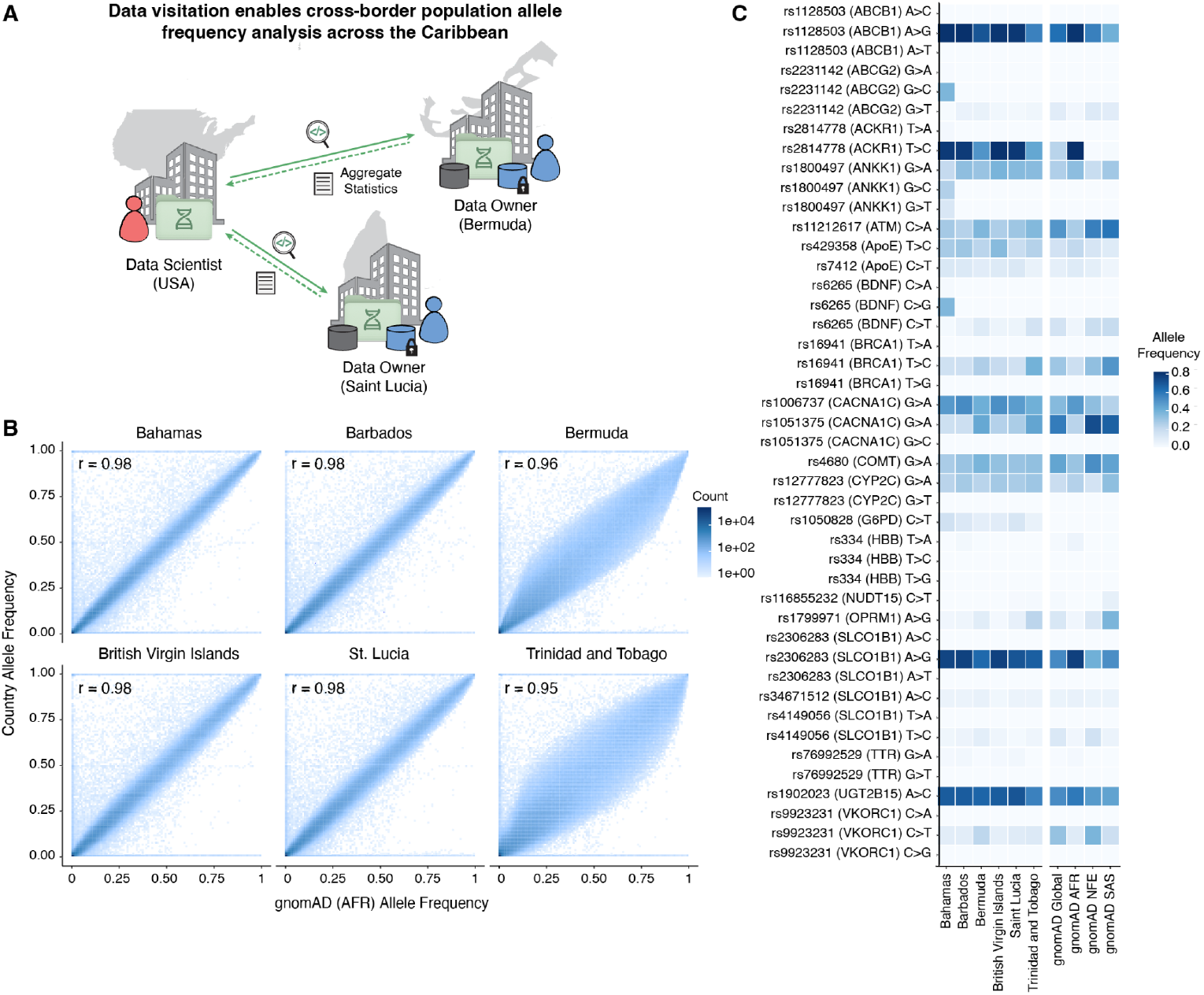
Cross-border analysis of population allele frequencies across Caribbean cohorts using data visitation. **(A)** Schematic of the BioVault-enabled remote allele frequency analysis workflow initiated from the United States and executed remotely at datasites in Saint Lucia (n = 97) and Bermuda (n = 488). Per-country SNP-array callsets from Caribbean cohorts, including the Bahamas (n = 168), Barbados (n = 125), the British Virgin Islands (n = 106), Bermuda (n = 488), Saint Lucia (n = 97), and Trinidad and Tobago (n = 76), were analyzed locally, and only aggregate allele frequency statistics were returned. **(B)** Two-dimensional binned scatter plot (100 bins) comparing per-country allele frequencies from Caribbean cohorts to allele frequencies from the gnomAD African ancestry reference population (AFR). Each bin represents the density of variants within the corresponding allele frequency range. **(C)** Heatmap of allele frequencies for a curated panel of pharmacogenetically and clinically relevant variants across Caribbean cohorts, displayed alongside reference frequencies from gnomAD global, African ancestry (AFR), non-Finnish European (NFE), and South Asian (SAS) populations.

Genome-wide allele frequency profiles from each cohort were compared with reference populations in gnomAD^44^ (**Fig. 4B, Supplementary Fig. 3**). Across all cohorts analyzed, allele frequencies were most strongly correlated with the gnomAD African ancestry (AFR) panel (Pearson *r* > 0.95 for all country-level comparisons), consistent with predominant African ancestral contributions to Caribbean populations. Bermuda and Trinidad and Tobago show the largest deviations from gnomAD AFR, reflecting greater admixture from non-African ancestry populations. This is consistent with reported population composition in these regions, including a substantial proportion of individuals of European descent in Bermuda (∼31%) and a highly heterogeneous population structure in Trinidad and Tobago, with large South Asian (∼35%) and African (∼34%) ancestry components alongside mixed-ancestry groups^34,35^.

To characterize population-specific patterns at clinically relevant loci, we analyzed per-country SNP-array callsets focusing on a curated panel of variants with established pharmacogenetic and clinical relevance (**Fig. 4C**). Country-level allele frequencies were calculated locally and compared with both global and ancestry-specific gnomAD reference panels. This analysis revealed marked heterogeneity in allele frequencies across Caribbean cohorts, with multiple variants showing population-specific enrichment or depletion relative to commonly used reference datasets.

Relative to gnomAD global allele frequencies, the most pronounced and consistent enrichment across Caribbean cohorts was observed at ACKR1 rs2814778 (T>C), which showed elevated allele frequencies in all populations analyzed, including the British Virgin Islands (AF 0.814, ΔAF +0.576), Saint Lucia (0.794, ΔAF +0.555), the Bahamas (0.788, ΔAF +0.549), Barbados (0.784, ΔAF +0.546), Bermuda (0.496, ΔAF +0.258), and Trinidad and Tobago (0.434, ΔAF +0.196). This variant is a well-established marker of African ancestry and is clinically relevant for hematological traits, including neutrophil counts and blood group phenotyping^36,37^. With varying magnitudes of enrichment among the islands, additional enrichments were observed at pharmacogenetically important loci, such as at ABCG2 rs2231142, which influences urate transport and response to drugs, such as allopurinol^38^, as well as SLCO1B1 rs2306283, which affects hepatic uptake of statins and other commonly prescribed medications^39^.

In contrast, several variants showed depletion relative to gnomAD global allele frequencies, including CACNA1C rs1051375, which has been associated with cardiac and neuropsychiatric phenotypes^40^, ATM rs11212617, linked to glycaemic response to metformin^41^, and VKORC1 rs9923231, a major determinant of warfarin dose requirements^42^. Bermuda and Trinidad and Tobago showed distinct patterns within the panel of clinically relevant variants, with Bermuda differing from other cohorts at loci including ACKR1 rs2814778, UGT2B15 rs1902023, and CACNA1C rs1051375. Trinidad and Tobago also showed divergence at variants including ACKR1 rs2814778, SLCO1B1 rs2306283, and VKORC1 rs9923231.

Together, these results demonstrate that remote allele frequency analysis via data visitation recapitulates known population structure and reveals substantial heterogeneity at clinically relevant loci across Caribbean cohorts, highlighting the importance of population-matched reference data and the utility of cross-border genomic analysis without centralizing sensitive genotype data.

### BioVault supports the design and deployment of secure computation frameworks for federated analyses

Federated computation across multiple datasites requires exchanging intermediate results during multi-step analyses. Sharing this data in plaintext, however, raises significant privacy concerns. Prior work has shown that even aggregate statistics of genotype data, such as allele counts or allele frequencies, can enable reconstruction or membership inference attacks^45,46^.

These risks are amplified in studies involving cohorts with modest sample sizes, fine-scale population structure, or enrichment for population-specific variants, where summary statistics may carry sufficient signal to enable re-identification or reveal community participation.

Cryptographic tools for secure computation offer promising strategies to address these privacy concerns in collaborative genomic studies^8,47,48^. However, designing cryptographic protocols for biomedical analyses over federated datasets requires deep expertise in both cryptography and distributed algorithms, as well as domain-specific knowledge in the target scientific domain^43^. Deploying these protocols across robust, decentralized networks requires a significant engineering burden^2,49^. Combined, these two factors limit the design and adoption of secure, federated solutions in the biomedical space.

To lower these barriers, BioVault utilizes Syqure – a custom-built Rust-based wrapper around Sequre^31^/Shechi^32^, a high-performance framework that translates Python-syntax pipelines into optimized, secure multiparty (MPC) protocols, homomorphic encryption (HE), and multiparty homomorphic encryption (MHE) using Codon. Syqure additionally provides a zero-configuration WebRTC proxy that establishes peer-to-peer connections between datasites behind firewalls, removing the requirement for publicly addressable TCP/IP network endpoints. This enables participants without a deep expertise in MPC/MHE frameworks to define and deploy custom secure computation protocols for biomedical workflows directly within BioVault.

As a proof-of-concept, we implemented a secure, federated protocol for calculating joint allele frequencies across Caribbean populations using BioVault’s pipeline framework, without requiring sharing of site-specific allele counts or sample sizes in plaintext (**Fig. 5A**). Each datasite locally aligned allele count vectors to a shared variant index (100k SNPs) and computed per-site allele counts and total allele numbers. These values were then split into encrypted shares and securely aggregated – coordinated across three BioVault desktop instances: two datasites holding private data and one aggregator serving as the trusted dealer (Methods). Secure element-wise addition produced pooled allele counts and total allele numbers, from which allele frequencies were computed by secure division. At no point during the protocol did any party gain access to another site’s per-variant allele counts, sample sizes, or allele frequencies; only the final pooled totals were revealed upon reconstruction of encrypted shares.

**Figure 5:**
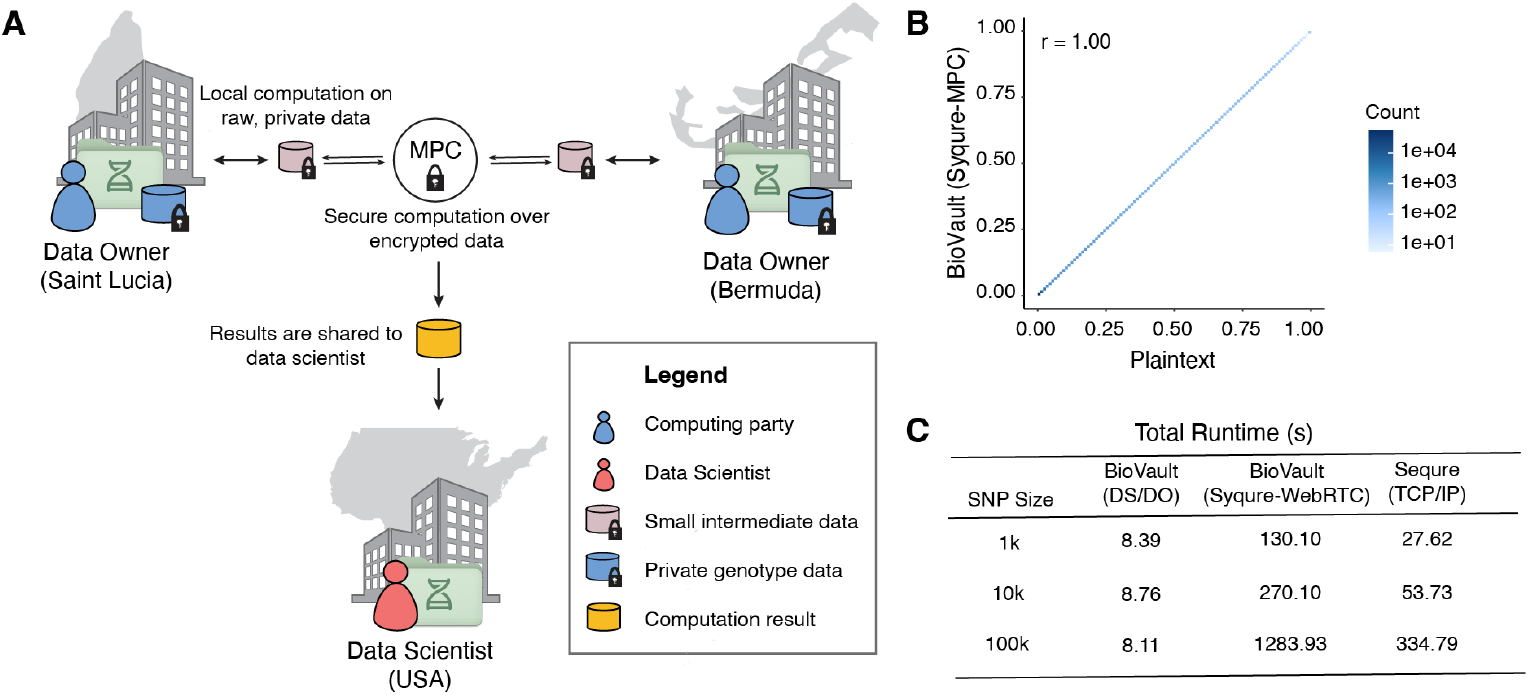
Secure federated allele frequency estimation across Caribbean datasites using multiparty computation. (A) Schematic of the BioVault-enabled secure multiparty computation (MPC) workflow for federated allele frequency estimation. Data owners (Saint Lucia and Bermuda) each perform local computation on private genotype data, including alignment to a shared variant index and calculation of per-site allele counts and numbers. Intermediate, per-site data are transformed into encrypted shares for downstream secure aggregation and secure division, coordinated across three BioVault desktop instances – data owners and one aggregator (remote server) serving as the trusted dealer. Only the final, joint allele frequency estimates are revealed; per-site allele counts and numbers remain encrypted and private. Data owners then shared the final estimates with collaborators in the United States. (B) Comparison of estimated allele frequencies obtained via our secure, federated computation on BioVault’s network and plaintext computation from raw allele counts and numbers. Concordance is exact (Pearson *r* = 1.00), confirming that secure aggregation introduces no numerical deviation from the plaintext reference. (C) Runtime benchmarks for secure, federated allele frequency estimation at three variant set sizes (1k, 10k, and 100k SNPs) across three execution configurations: (1) BioVault-enabled data visitation paradigm without MPC (DS/DO), (2) BioVault with Syqure (Sequre-MPC over WebRTC), (3) Sequre MPC over direct TCP/IP. WebRTC-mediated MPC, as implemented by Syqure, introduces approximately a 4–5x overhead relative to direct the TCP/IP network connections implemented in Sequre. This slowdown is likely attributable to relay-assisted NAT traversal and connection negotiation. However, this overhead enables deployment across firewalled institutional networks without requiring publicly addressable endpoints. In future work, we aim to address this added latency from WebRTC to minimize the gap in performance.

As expected, the resulting joint allele frequency estimates were exactly concordant with those obtained from plaintext computation based on access to raw allele counts and numbers (Pearson *r* = 1.00; **Fig. 5B**). To assess scalability of BioVault’s network, we assessed runtime of our secure, federated workflow across three variant set sizes (1k, 10k, and 100k SNPs) across three execution configurations: (1) BioVault-enabled data visitation without MPC, (2) BioVault with Syqure (Sequre-MPC over WebRTC), (3) MPC over direct TCP/IP as implemented in Sequre/Shechi. BioVault’s plaintext data visitation workflow, without cryptographic integration, completed aggregation in under 10 seconds at all scales (**Fig. 5C**). Execution of the same computation in a privacy-preserving manner with Syqure introduced an additional 4–5x overhead relative to out-of-the-box Sequre. This is likely attributable to relay-assisted NAT traversal and connection negotiation required by BioVault’s WebRTC network, compared to Sequre’s TCP/IP protocol. Despite this overhead, BioVault’s WebRTC-mediated deployment enables communication across firewalled institutional networks, without the need for without the need for complex firewall reconfigurations, static IP assignments, or manual port forwarding.

These results demonstrate BioVault’s utility in bridging the gap between the design of secure computation protocols for biomedical analyses facilitated by the integration of cutting-edge MHE/MPC compilers like Sequre/Shechi, and their deployment across real-world computing environments.

## Discussion

Here, we presented BioVault, an open-source, privacy-first, data visitation platform for global collaboration in biomedicine. BioVault enables remote analysis without raw data transfer, while preserving analytical fidelity and institutional control. Designed for out-of-the-box use, BioVault lowers technical barriers that have historically limited meaningful biomedical collaboration to well-resourced institutions. This supports inclusion of emerging biobanks, LMIC institutions, and Indigenous communities as analytical partners rather than data contributors alone – creating more equitable partnerships.

We showcased BioVault’s versatility and adaptability by performing remote analysis on various data types and structures, such as scRNA-seq, physiology, MRI, and whole genome sequencing data. We further demonstrated BioVault’s utility in enabling cross-border collaboration through two remote analyses: a genome-wide association study of Type II Diabetes in Jordan’s Circassian and Chechen populations and population allele frequency estimation across Caribbean cohorts. We also showed that BioVault can lower the barrier to adoption of cutting-edge secure computation frameworks in the biomedical space through a secure, federated estimation of joint allele frequencies across the Caribbean, using MPC.

BioVault’s data visitation platform adopts a governance-based system that emphasizes transparency, auditability, and explicit authorization of computations over private, locally held data. Every execution request is logged, auditable, and subject to human approval before results are released. Pipeline development based on our “twin” data paradigm using mock, synthetic data ensures that workflows can be validated without exposure to real data, and encrypted transport of intermediate results further minimizes privacy leakage. This governance-based approach, however, depends on careful review of execution requests and responsible analytical design. Malicious or poorly constructed workflows, dependency-level exploits, repeated querying, or structured outputs may enable information leakage. Open challenges include enforcing global privacy budgets across repeated analyses and verifying that executed code and returned outputs faithfully match approved workflows. Addressing these risks will require technical safeguards such as privacy-budget accounting^50^, zero-knowledge proofs^51–53^, and hash-based verification of model inference^54^.

Acknowledgement of these privacy risks motivated our integration of secure computation frameworks, based on modern cryptography, which offer stricter and more formal privacy guarantees. These frameworks are especially important in joint computations across federated datasets, where raw data is locally held but shared intermediate outputs or aggregate summaries can still enable re-identification, particularly in small or underrepresented cohorts. To address this, BioVault implements Syqure by combining Sequre/Shechi, a high-performance compiler for MPC/MHE, within our network layer. Participants can design and deploy custom secure computational workflows across federated datasets, with little to no configuration through our desktop application. While this capability is powerful, it remains under active development.

Future work will focus on improving scalability, strengthening safeguards against cumulative leakage across repeated analyses, and enhancing interoperability.

Future development will also emphasize interoperability with emerging international standards for federated genomic data sharing, including those advanced by the Global Alliance for Genomics and Health (GA4GH)^55^. Although BioVault enables remote analysis of sensitive data, its deployment must remain aligned with existing legal, ethical, and policy frameworks. Use of the platform requires compliance with data access agreements, regulatory requirements, IRB oversight, and informed consent policies. As technical capabilities evolve, clarification may be needed regarding whether remote execution and derived outputs constitute “data sharing.” By operating within established governance structures rather than circumventing them, BioVault supports privacy-preserving collaboration while reinforcing institutional accountability and regulatory compliance^56–58^.

As large-scale initiatives such as the Human Genome Project II^59^, the Human Immunome Project^60^, and the Human Phenotype Project^61^ expand the scale and diversity of genomic and health-related data, enabling these datasets to be analyzed collectively, while respecting sovereignty, regulation, and governance, will be critical for advancing human health. Although the value of health data increases with aggregation, cross-border transfer restrictions and the burden of centralization often limit collaboration – particularly for under-resourced institutions. By decoupling analysis from data movement, BioVault enables distributed, interoperable workflows across sites while each institution retains custody and oversight. Through data visitation, compatibility with secure computation frameworks, and a low barrier to adoption, BioVault will provide a foundation for scalable and inclusive biomedical collaboration as global data generation accelerates.

## Figures

**Supplementary Figure 1:**
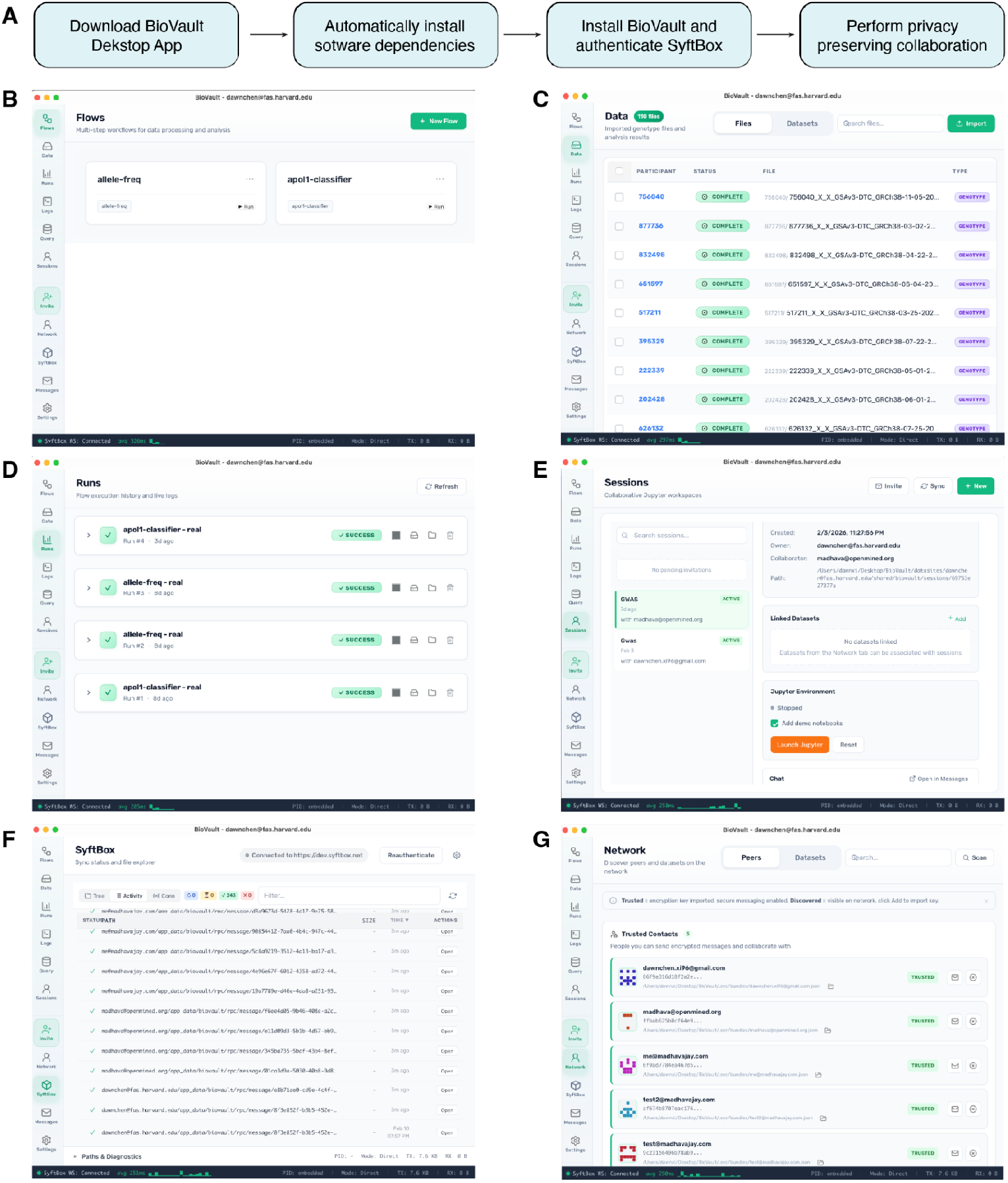
Overview of the BioVault Desktop application. **(A)** Schematic of the installation workflow for the BioVault Desktop application, illustrating the streamlined setup process designed to facilitate ease of use. **(B–F)** Representative screenshots of the BioVault interface: (B) Flows page for configuring and managing computational workflows; (C) Data loading interface; (D) Flow results page displaying analysis outputs; (E) Interface for initiating Jupyter-based analysis sessions; (F) SyftBox networking page; (G) Visualization of the BioVault user network.

**Supplementary Figure 2:**
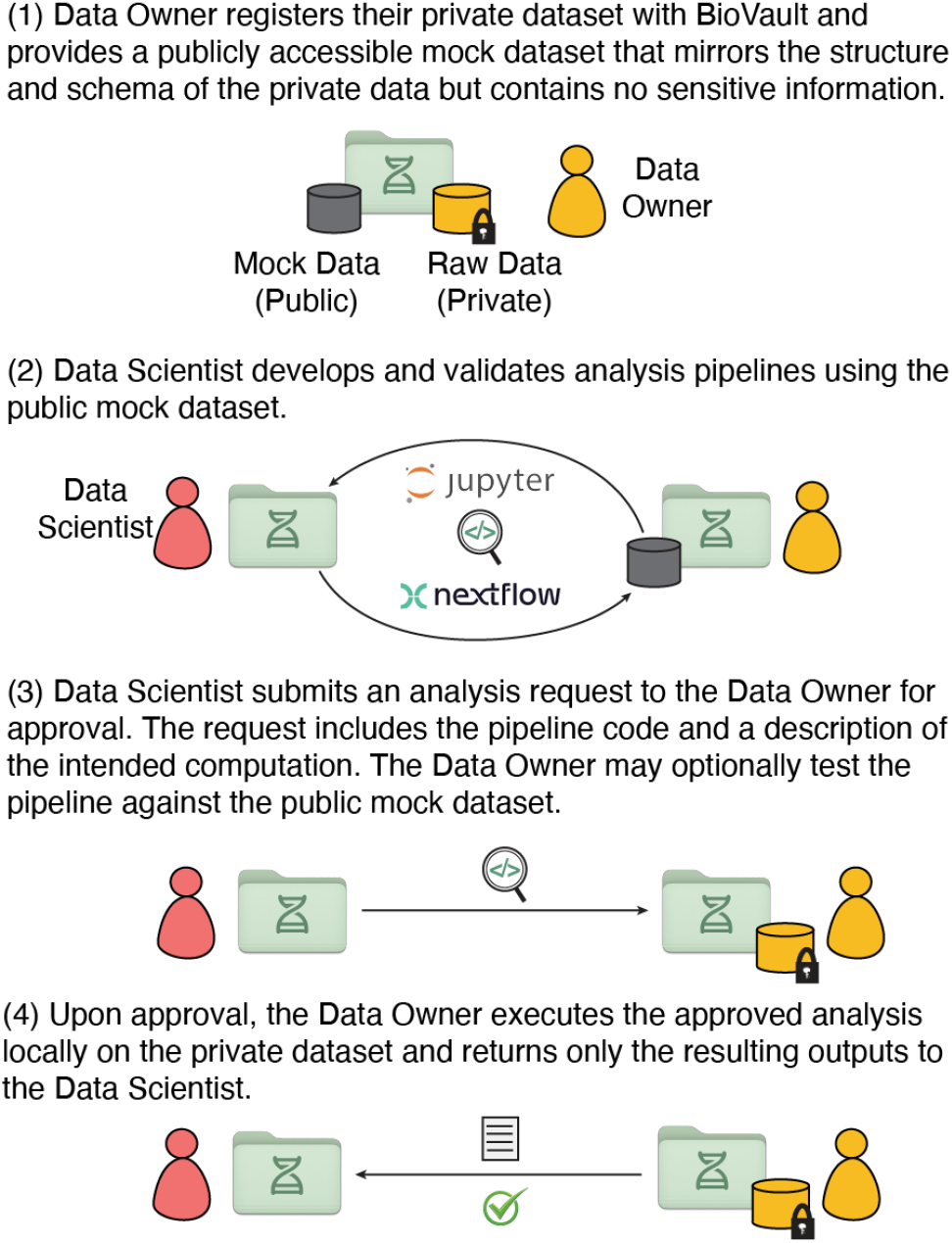
BioVault “twin” paradigm for privacy-first, iterative pipeline development and execution. Data owners publish a synthetic dataset that mirrors the schema of a corresponding private dataset without exposing sensitive content. Researchers develop and validate analysis pipelines against this synthetic twin, then submit execution requests via the SyftBox network. Data owners review and authorize requests under local governance policies. Approved computations are executed within the data owner’s environment, with complete audit logging, and only explicitly permitted outputs are returned. This decentralized, attribution-based model enables analysis across heterogeneous compute environments without data centralization while preserving institutional control and data sovereignty.

**Supplementary Figure 3:**
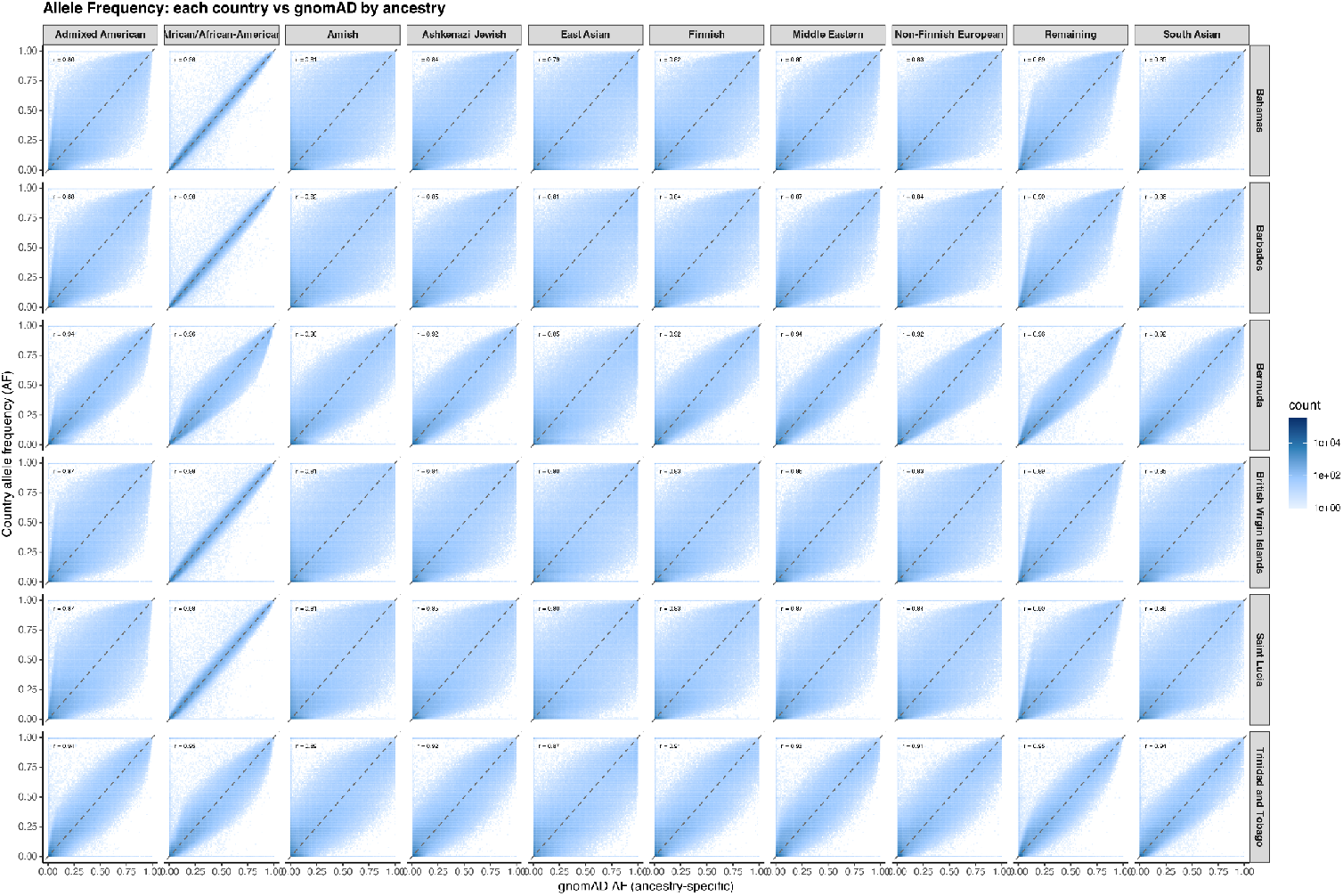
Cross-Population Comparison of Allele Frequencies Between gnomAD and Caribbean Cohorts. Two-dimensional binned scatter plot (100 bins) comparing per-country allele frequencies from Caribbean cohorts to allele frequencies from gnomAD across all ancestry classes. Each bin represents the density of variants within the corresponding allele frequency range.

## Materials and Methods

### Biovault Architecture

BioVault is an open-source, modular software framework that operates over an end-to-end encrypted, peer-to-peer (P2P) network with optional relay-assisted connectivity. The system is composed of interoperable components implemented across multiple languages to balance performance, portability, and ecosystem integration.

The BioVault ecosystem includes the following primary packages that interoperate:

- BioVault CLI (Rust)
- BioVault Desktop (Rust / JavaScript)
- BioVault Beaver (Rust / Python)
- SyftBox (Go / Rust)
- SyftBox SDK (Rust / Python)
- SyftBox Crypto (Rust)
- Syqure (Rust / C++ / Codon)

The BioVault command-line interface (CLI) and core libraries are written in Rust to ensure high performance, memory safety, and broad platform compatibility. Rust’s toolchain enables compilation for macOS, Windows, and Linux across major CPU architectures.

BioVault relies on the SyftBox SDK for networking and cryptographic functionality. SyftBox provides end-to-end encryption, byte transfer, file synchronization, and peer discovery across distributed nodes. A reference relay server implementation is written in Go and leverages object (blob) storage for caching and message exchange, with support for Amazon Web Services (AWS) S3 and S3-compatible systems such as MinIO or SeaweedFS. SyftBox clients, implemented in Go and Rust, authenticate with the relay server and synchronize encrypted file trees between local systems and remote collaborators.

### System Components and Execution Model

BioVault CLI and BioVault Desktop enable users to register private datasets and optionally publish synthetic or mock datasets for workflow development. Remote collaborators can construct and submit computational workflows that execute against the hosted data under a data visitation model.

Encrypted communication, file transfer, and firewall/NAT traversal are handled by SyftBox through its SDK and cryptographic libraries. All data transfers are end-to-end encrypted.

BioVault orchestrates computational workflows using a declarative YAML-based specification termed a Flow. Flows define execution steps and can invoke external workflow engines (e.g., Nextflow), shell scripts (bash), or privacy-preserving computation frameworks, including secure multi-party computation (SMPC) and homomorphic encryption (HE) via the Syqure/Codon compiler.

BioVault Beaver provides an interactive Python-based Jupyter notebook interface that supports exploratory and iterative execution within the visitation framework.

The BioVault Desktop application integrates these components into a graphical user interface compatible with macOS, Windows, and Linux. The desktop application includes pre-bundled binary dependencies, dataset search and discovery tools, data import and metadata (dataset card) creation workflows, job history tracking, a local SQLite database for result recording, and an end-to-end encrypted messaging interface supporting one-to-one and group communication.

### Connectivity, Identity, and Addressability

SyftBox supports two complementary communication modes: asynchronous file synchronization and real-time peer-to-peer (P2P) networking.

The majority of SyftBox operations occur over an asynchronous file synchronization layer that tolerates intermittent connectivity and can operate in offline-first environments. Data may be synchronized via the filesystem and relayed when network connectivity becomes available, making this mode suitable for personal computers and heavily firewalled institutional systems. This layer functions primarily as a byte-transfer and state synchronization mechanism that is agnostic to higher-level encryption and identity abstractions. It enables modeling of a globally addressable virtual file path for private datasets and enforces Unix-style permission controls supporting multi-owner read, write, and administrative access. Relay infrastructure accelerates the transfer of large files, including models and datasets (real or synthetic), while maintaining encrypted transport.

For network-intensive computations requiring low latency—such as secure multi-party computation (MPC) or homomorphic encryption (HE)—SyftBox supports real-time encrypted communication using WebRTC-based peer-to-peer connections, facilitated by a TURN server for NAT and firewall traversal. Although such computations can be executed over the asynchronous synchronization layer, performance is substantially improved through direct P2P connectivity.

To enable decentralized resource referencing, SyftBox implements the syft:// protocol. Unlike conventional http://addressing—which depends on centralized domain name resolution (DNS), certificate authorities for TLS, and continuously reachable public IP infrastructure—the syft:// scheme references resources by datasite identity and path, with resolution handled through relay-assisted peer discovery. In the current implementation, a reference relay server is operated centrally; however, the protocol is designed to support multiple independent relay operators, enabling federated or decentralized deployment models.

Identity is currently implemented using an email-based registration system, reflecting its widespread use and established trust model in scientific collaboration. Users register by verifying control of an email address through a two-factor authentication (2FA) challenge, after which a web authentication token is issued for server interaction. Although email serves as the initial identity mechanism, the protocol is identity-agnostic by design and supports future integration of alternative identity systems, including privacy-preserving or decentralized identifiers. Notably all encrypted files contain signatures from a users offline private key which is unrelated to the email web token auth flow.

End-to-end encryption is supported through user-published public key material. Each user may upload a did.json file containing cryptographic public keys at a well-known path, enabling signature verification and encrypted communication between peers. While optional at the protocol level, BioVault enables this functionality by default to ensure encrypted data exchange and authenticated execution.

An example syft:// url for a public / private resource would be:

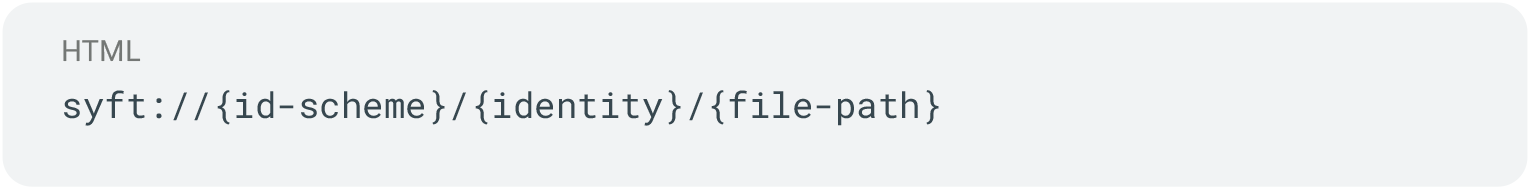

For email we have defaulted to no {id-scheme} for now, like so:

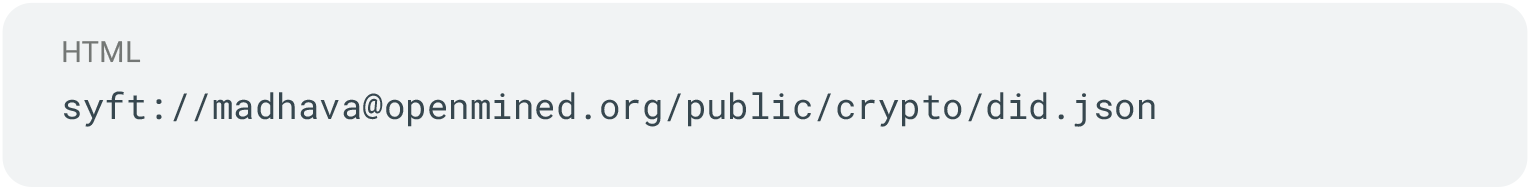

This is optionally also made available over HTTP via the cache server via:

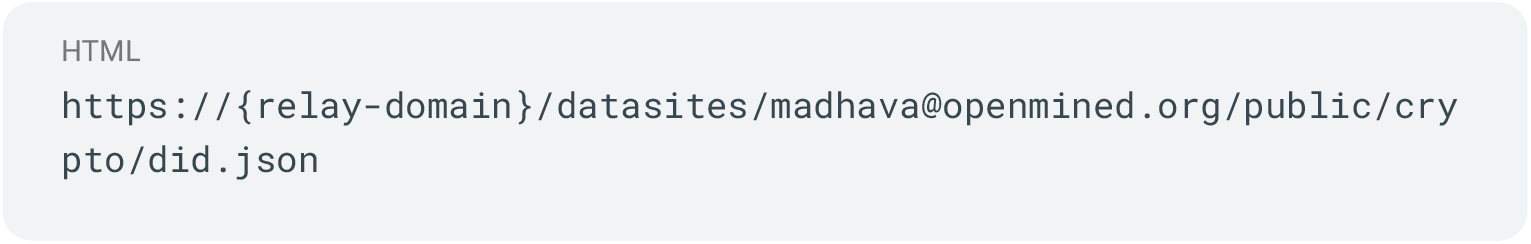

Note the file path stub is identical. This syft:// path is canonicalized from a datasite root on the users machine preventing path traversal referencing.

To address private resources, we provide an alternative path that does not actually exist.

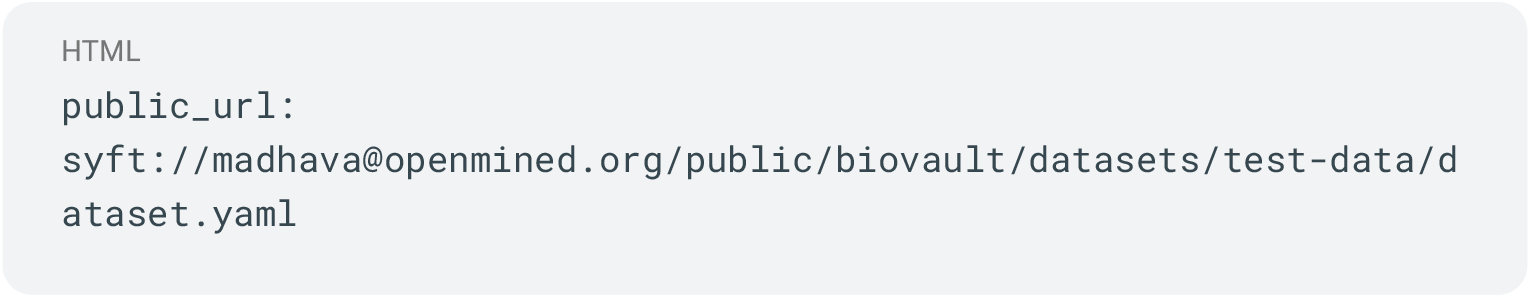
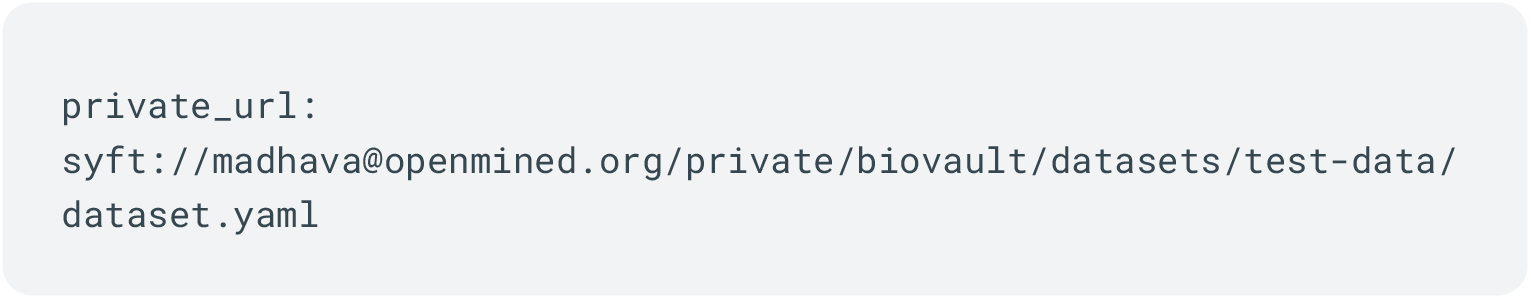

When linking data into BioVault we create a mapping between its real location and this canonical syft url which is not made available to syftbox or other users.

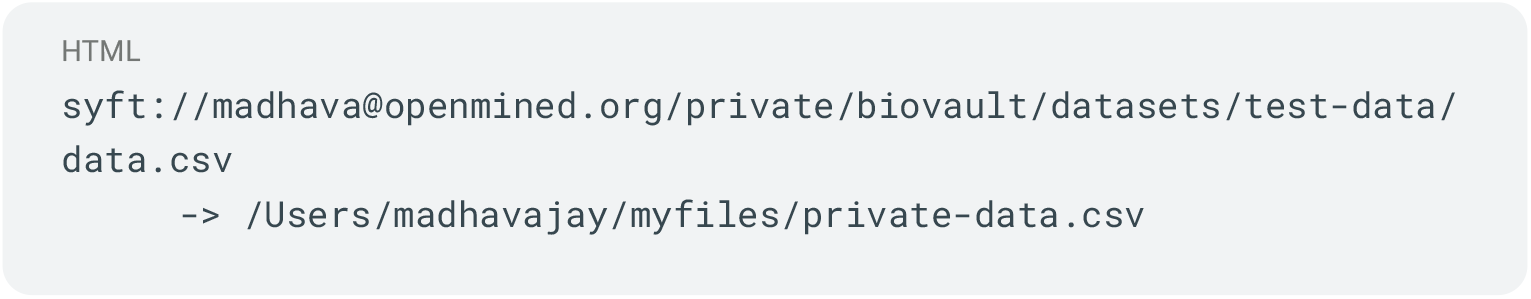

This file is kept entirely out of the system and can be used to resolve private urls at run time.

In this way, public or mock data can be accessed directly via https://orsyft://via a relay server to download locally. These local mocks can be used to construct computations and execution can be requested against the real private data by binding it to this unique url.

To allow both human and machine interpretability we publish a yaml specification for datasets which can optionally include mock / sample data.

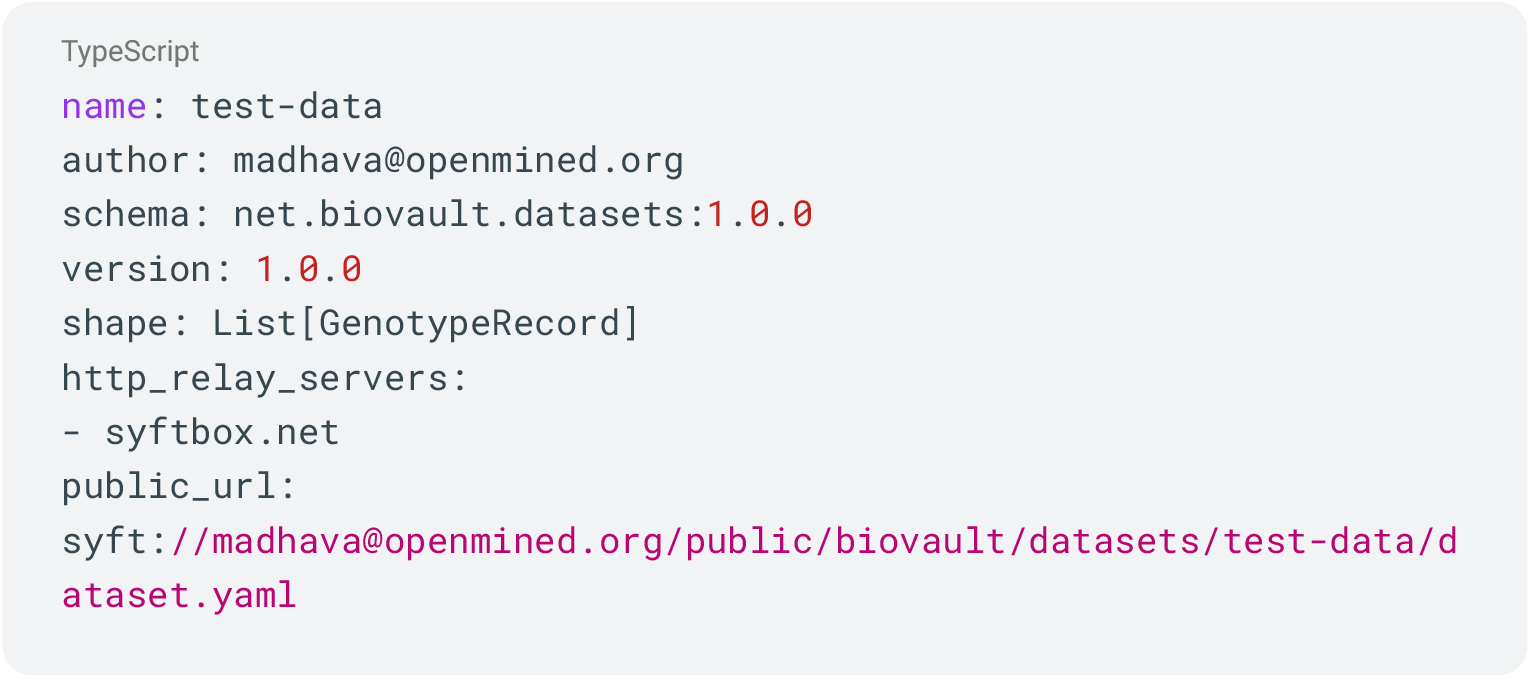
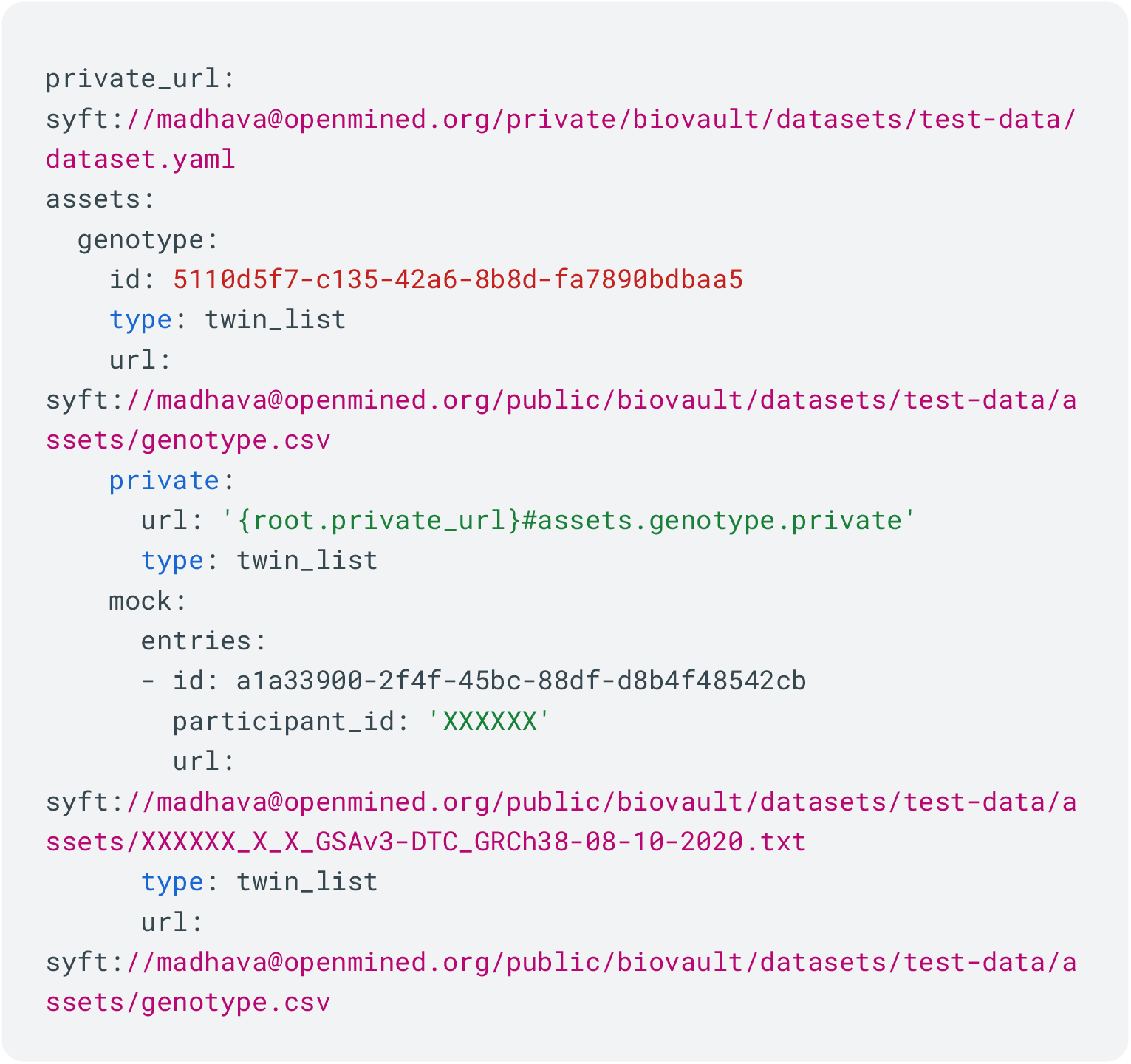

There are several record types including lists and dictionaries and support for the Twin paradigm. Consuming code can operate on a copy of the mock path and then bind to the private when requesting computation on the source data.

### Privacy

BioVault operates over the SyftBox peer-to-peer synchronization layer, which provides authenticated, end-to-end encryption of messages and files without being online. Confidentiality and integrity of user data are enforced at the application layer prior to any network transmission. Traffic to the server is delivered over HTTPS and websockets or WebRTC.

Data are encrypted locally using the recipient’s public key and written atomically into a SyftBox synchronization path. The underlying byte-transfer mechanism operates on ciphertext and does not access plaintext data except for opt-in public metadata. Plaintext is decrypted only into a separate local path after transfer and is never exposed within the synchronization subsystem.

For low latency connections such as during SMPC / HE operations with Sequre we utilize WebRTC using rust library webrtc-rs which provides DTLS 1.2.

BioVault uses Ed25519 keypairs for identity and digital signatures and X25519 for key agreement. Pairwise authenticated encryption is established using an Extended Triple Diffie–Hellman (X3DH) protocol and ChaCha20-Poly1305 as the authenticated encryption with associated data (AEAD) scheme.

Security relies on the hardness of the elliptic-curve discrete logarithm problem over Curve25519 and the IND-CCA2 security of ChaCha20-Poly1305. The implementation leverages the open source rust libraries ed25519-dalek, x25519-dalek, and chacha20poly1305.

This provides the following guarantees:

- **Confidentiality** (IND-CCA security): ciphertexts reveal no information about plaintext to passive or active adversaries without access to the corresponding private key.
- **Integrity and authenticity** (EUF-CMA security): modifications to ciphertext, public key changes or impersonation attempts are detected.

Forward secrecy via continuous ratcheting and Post-quantum cryptographic resistance are not provided at this stage but would make for good follow up enhancements.

Additionally zero knowledge proof identity mechanisms would be a welcome addition.

BioVault and Syftbox use a TOFU (trust on first use) model. Users should verify public keys out of band and monitor for changes.

Biovault warns users if the key they have for someone has changed and requires them to opt into trusting a replaced key which could be the sign of an attacker attempting to hijack the communication. To further help users do out-of-band verification, we display an easy-to-recognize bitmap image of their public key to visually compare when keys have changed.

The public key material is stored in the well-known path: syft://{email}/public/crypto/did.json in a did:web compatible json format.

The private key is stored in a folder outside the datasite sync path and a BIP39 (key phrase) recovery key is displayed to the user once during key creation and can be stored safely and easily using third party password management software.

When making data available to multiple recipients, we encrypt the files with each of their keys such that each recipient can also decrypt the file.

To make data available to another user of the system we write it to a folder with a syft.pub.yaml which describes what users can read or write to which paths with a flexible glob pattern-based system and unix style read, write, owner groups.

Example publish rules:

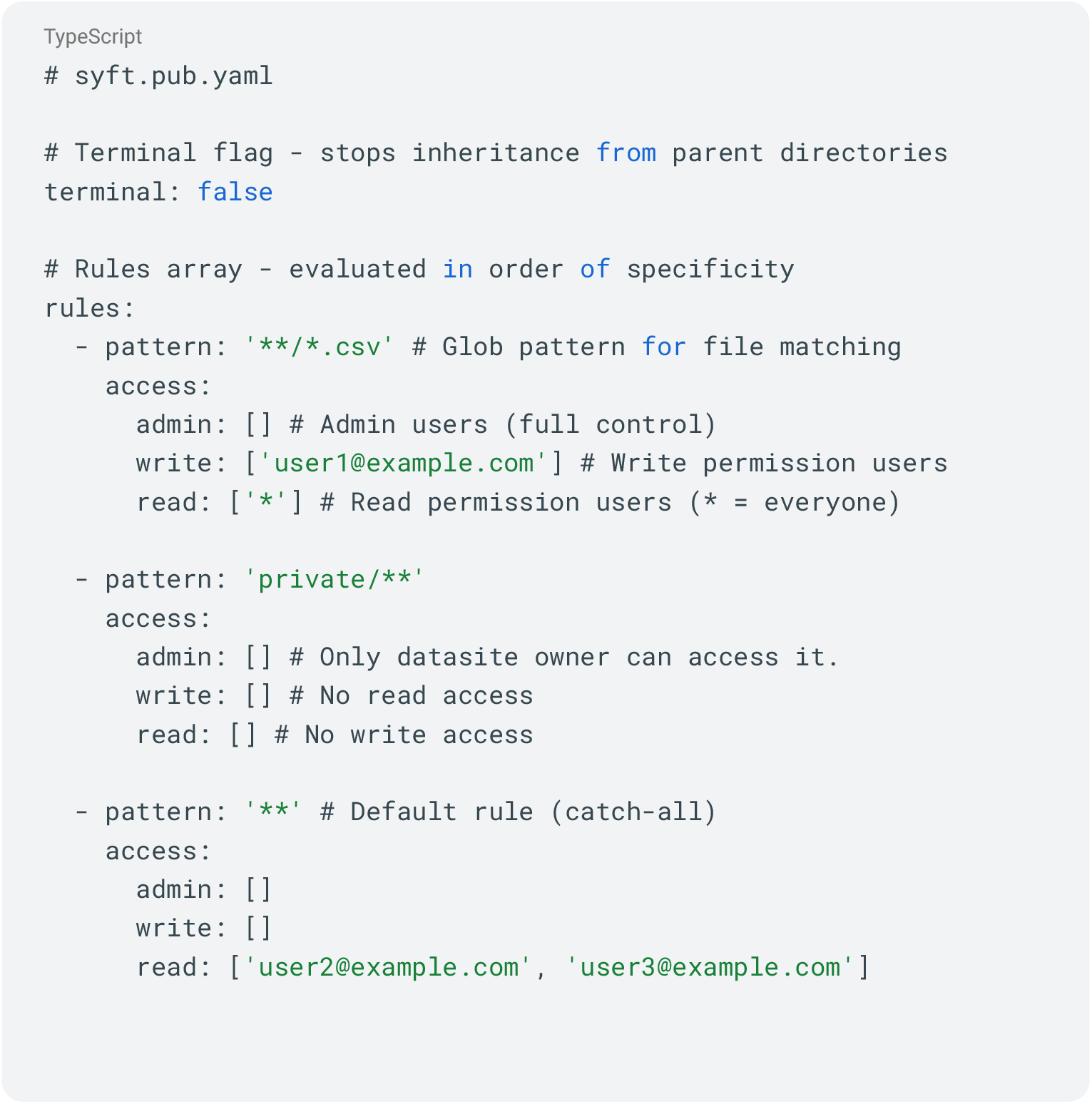

### Special Tokens

We provide special tokens to allow the construction of useful patterns.

- ‘*’: Wildcard representing all users (public access)
- ‘**’: Glob pattern matching all files recursively
- ‘USER’: Dynamic token that resolves to the requesting user’s ID

RPC messages which expect a response are sent via this same system to a pre-defined path which can use the USER token to automatically bucket messages by user.

Incoming messages are given a time sortable uuidv7 id and the.request extension and.responses are written back by the receiver next to them.

Each user opts into synchronization using a syft.sub.yaml file. Data is only delivered when the recipient subscribes to a given path including public keys and RPC messages allowing customizable firewalling.

Many operations in BioVault such as creating sessions or shared computations creates rules for subscription for each party to a unique path on each datasite for said operation.

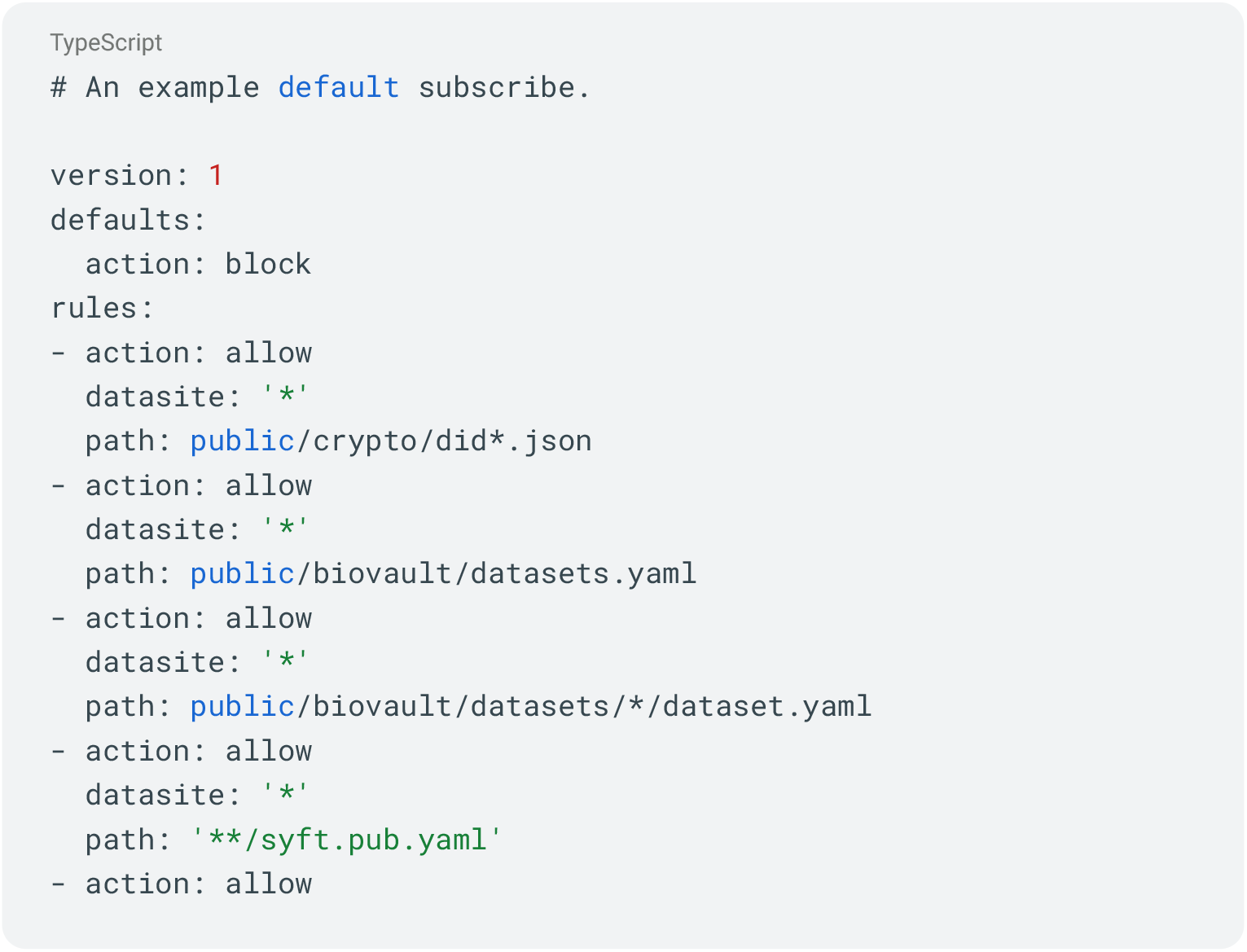
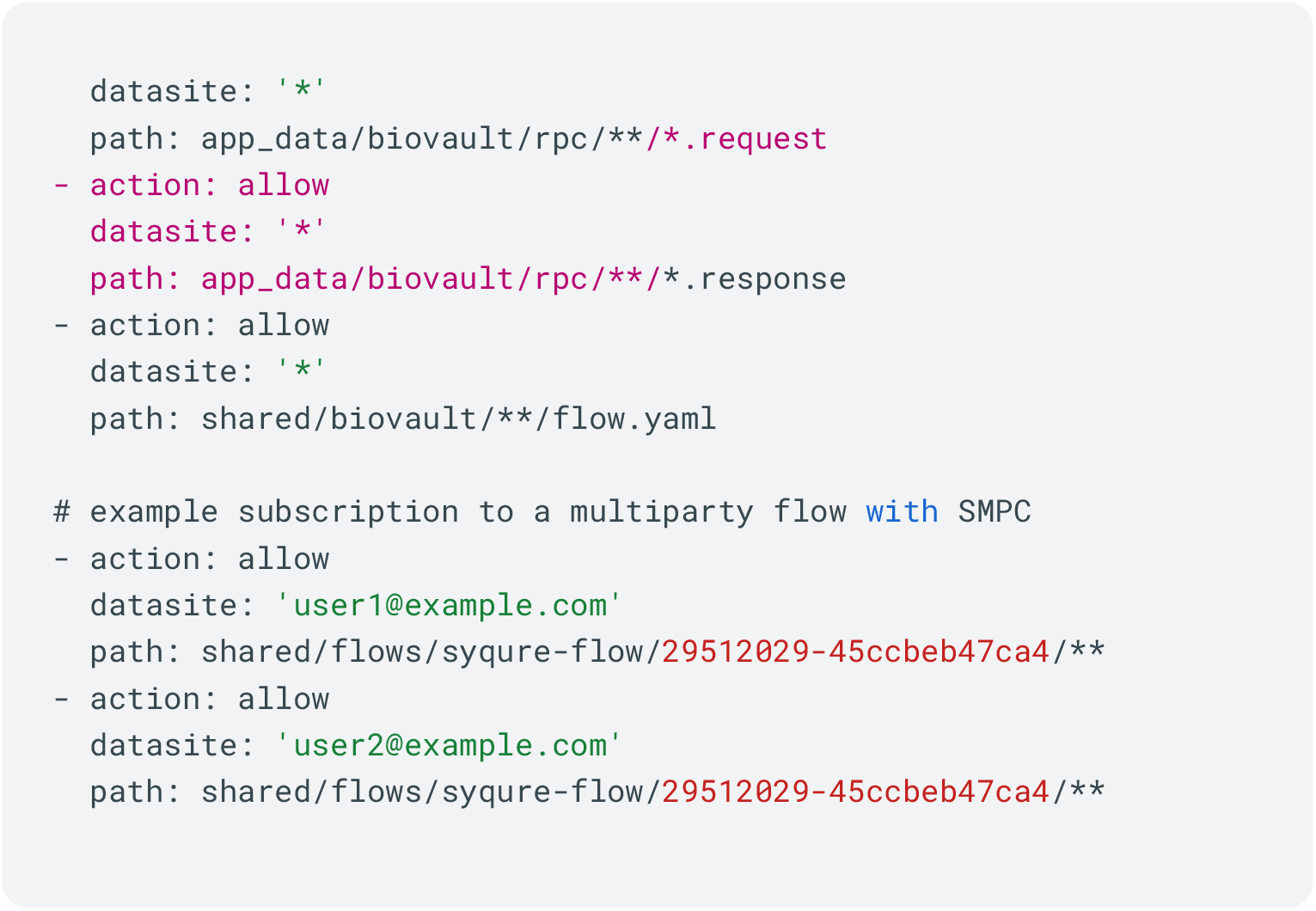

### Computation

We offer two key paradigms to produce shared computation, one which can be executed iteratively via a Jupyter notebook and python library and another which allows arbitrary pipelines of fixed computation to be constructed with one or more parties utilizing one or more backends such as Nextflow, Bash and Sequre.

### Dynamic Jupyter Eager Execution

To collaborate via an iterative Jupyter notebook, users can invite each other to a session which allows the launching of a managed Python and Jupyter interface on their local machine. Both parties have their own Jupyter instances and do not share access to their kernels or an active HTTP connection. All interaction is via serialized files that are encrypted and shared with each other over SyftBox.

The notebook interface includes some common libraries such as pandas and numpy and an additional library from biovault called biovault-beaver (eager execution) which facilitates a secure human-in-the-loop back and forth model for data exploration and data science.

In beaver both parties are equal peers so the distinction of who governs what data is arbitrary, however we include a number of example notebooks that utilize public data to allow users to test and reference the Data Owner / Data Scientist paradigm. While we didn’t build it there is no reason these sessions cannot facilitate n parties in the future.

Once logged into a session in Jupyter, communication is handled via a series of requests sent and received using a Python API.

Messages including shared data, computation requests and computational results are available in the bv.inbox(). These files are written from each side to a self-governed path in SyftBox that is made read-only to the other participant. Each person builds up their communication by writing to their side and the other reads from the opposing side.

In a similar manner to other parts of Biovault and SyftBox data is kept in two different locations. The Jupyter kernel runs at a path outside of the syftbox synchronization path.

For example:

/Users/username/Desktop/BioVault/sessions/7a3509ae19f5

While the synchronized data is shared via a path which SyftBox keeps synchronized:

/Users/username/Desktop/BioVault/datasites/{users-email}/shared/biovault/sessions/7a3509ae1 9f5

Beaver then knows to monitor the peers folder at:

/Users/username/Desktop/BioVault/datasites/{peers-email}/shared/biovault/sessions/7a3509ae1 9f5

We utilize fory (and pyfory) an apache serialization library which provides performant and platform agnostic data serialization for python objects (and other languages like R). In the event that objects require custom serialization steps we allow users to construct their own serializers using decorators and have those bundled and sent along with their data dynamically at run time.

For instance this code provides serde for scanpy / anndata

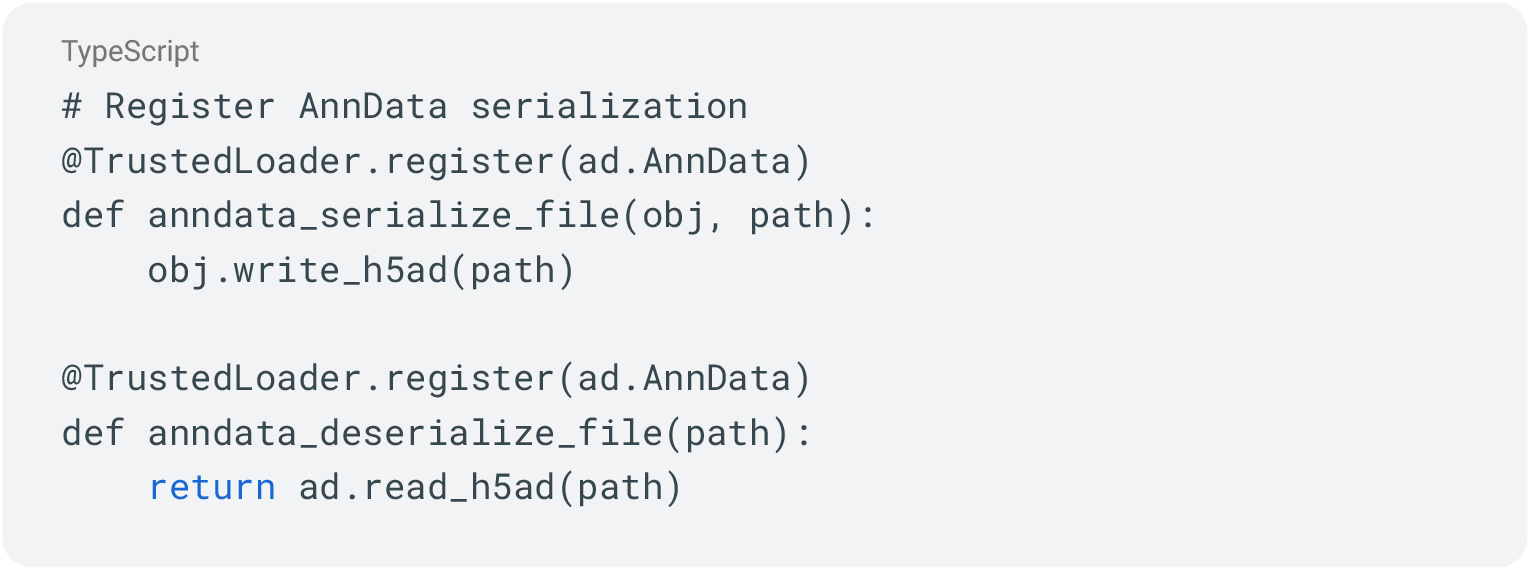

The API includes methods to test the roundtrip of your custom serializer making rapid development to support dynamic serde of arbitrary object types easy.

Since serialization of data can be an attack vector we avoid the use of pickle and ask users to review the plaintext version of custom deserialization code before executing it.

The python ecosystem is enormous, and while many libraries will work out of the box with pyfory, many will not without custom serializers. Future work would include providing a large set of verified and tested python library support.

Data can be made available to the other party as a Twin object either before the session or during the session only using the Python API.

Special python specific remote_vars can be made available to the other participant dynamically during execution of the session and read on the other side as though they were local objects, even allowing the assignment of global namespaced variables to provide a 1-1 experience when referencing objects. During loading we prompt users to opt in to these automatic variable assignments as it’s not always desirable.

Additionally, remote variables can be marked as live which auto publishes them when they change allowing the construction of useful remote_vars that contain metadata such as model training accuracy to be shared in real time as programs execute on private data remotely.

Biovault Beaver also includes syftbox-sdk via a maturin compiled rust binary to provide beaver with the ability to parse syft:// urls, read and write to syftbox paths and encrypt and decrypt files.

To construct and test computation in beaver, users simply need to decorate a normal python function with ‘@bv’ turning the function into an object like so:

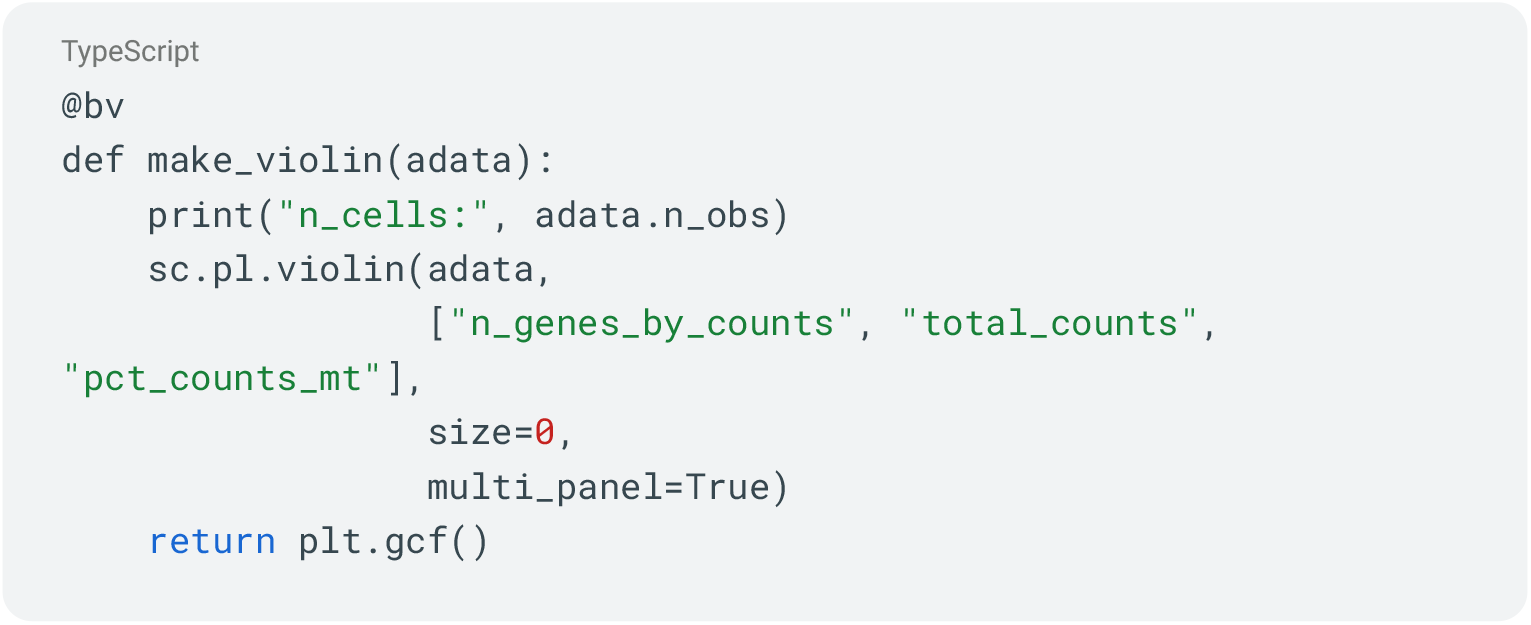

Once executed like a normal function on a Twin object obtained from remote_vars or the inbox, users can see what the result would be on mock data ensuring their computation matches the expected python object structure and logic.

The bv decorated function transparently deboxes the mock side of the twin when executing against it so that only the real python object interfaces with the real python code preventing wrapper object / monkey patching problems.

The return value is then boxed back up into a Twin.

Users can then request the other party run their code against the real data by using simple.syntax.

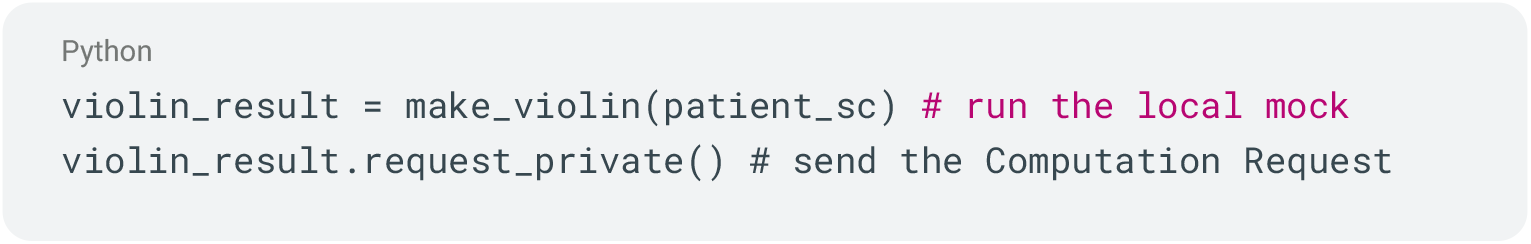

Data owners can receive these requests in their notebook, inspect them and the code they contain and then opt to run them against a mock of their twin to determine if they run as expected.

Once satisfied, data owners can execute the code against real data and then review it before choosing to release it back to the requester. Much of notebook based data science relies on emitting side effects within standard out (stdout), standard error (stderr) and plots during function execution to aid in debugging and visualizing code outputs. However these can also leak private information under remote execution flows.

Beaver captures these details and makes them available as twins, allowing data owners to review any prints or plots from private data and choose to withhold or release them.

Plots are captured as png to prevent references to underlying raw data being serialized and transmitted, but data owners can optionally extract the original plots from the object if required.

Once this returned result is made available the data scientist can load this from their inbox, choose to deserialize it and load its variables and the system will update any existing references to the original twin with the private side.

This means that the data scientist can now continue their notebook with the same Twin variable and all future computation will continue preferencing the private real result side now by default.

This pattern can continue on indefinitely either choosing to reference the original data or derived variables to arrive at either a final result or a series of computational steps which can be bundled up and saved for future use.

### Static Flows and Modules

In a similar fashion to above, Biovault provides a way to define upfront computation called Flows which can be executed collaboratively by sending them to users via the chat interface, url or file system.

Within the chat interface, users can sync the flows and then review them on disk. Once satisfied they can be run either against mock or real data and the results can be returned to the user via the same chat interface.

For multiparty computations, the chat interface supports multiple person groups and flows can be written to support more than one individual split into arbitrary roles.

Results from execution can be optionally captured in the local sqlite database for future reference providing audit trails; and follow up analysis can be performed using SQL directly within the query tab and displayed query tables exported to csv.

This flexible syntax allows for users to define arbitrary single or multiparty computations and share progress state and logs so that each party can see real time updates to the online computation.

Below is an abridged example of an example flow file with submodules that performed the secure aggregation of allele frequency summary files in our experiment using codon + sequre to perform an MPC with Additive Secret Sharing.

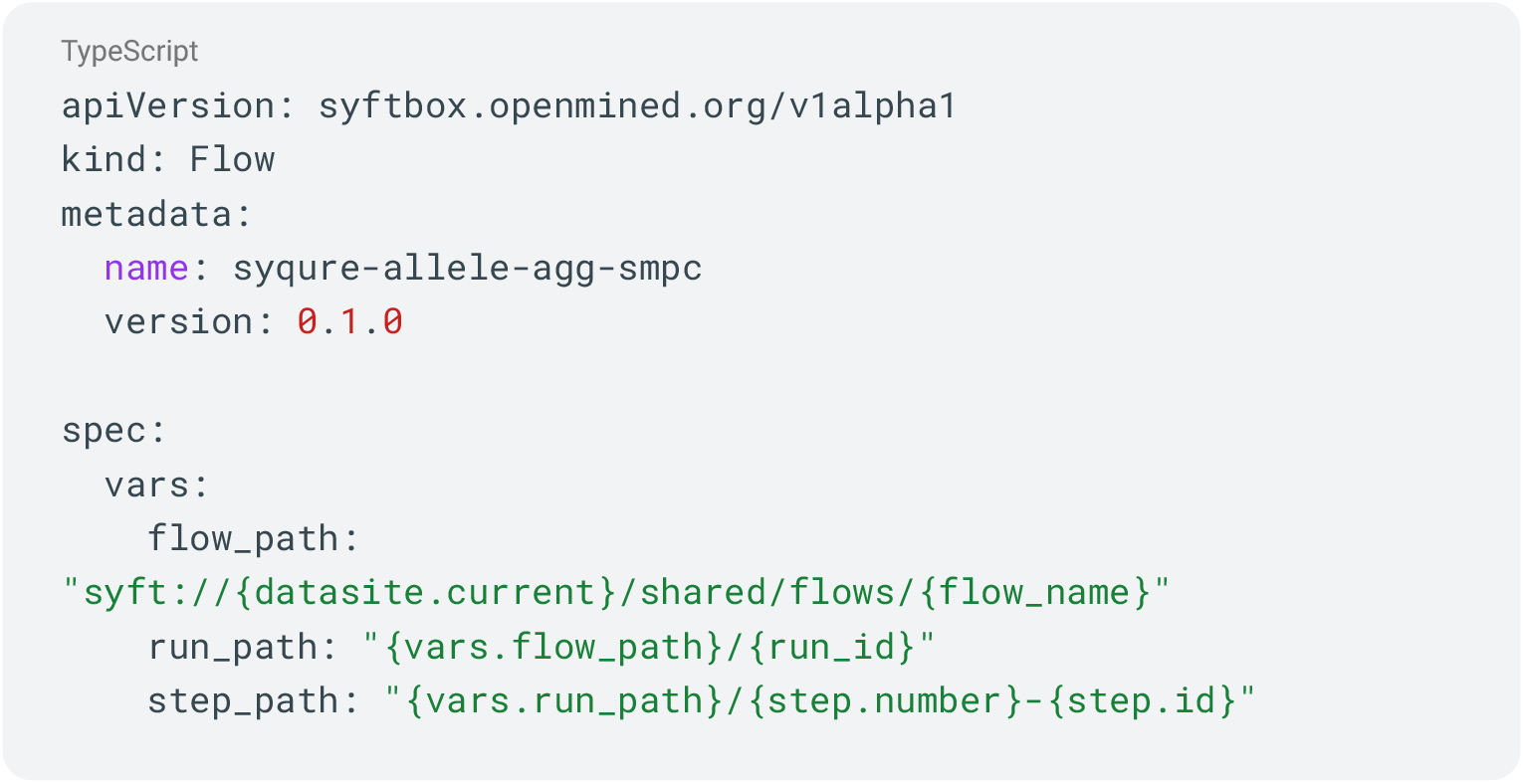
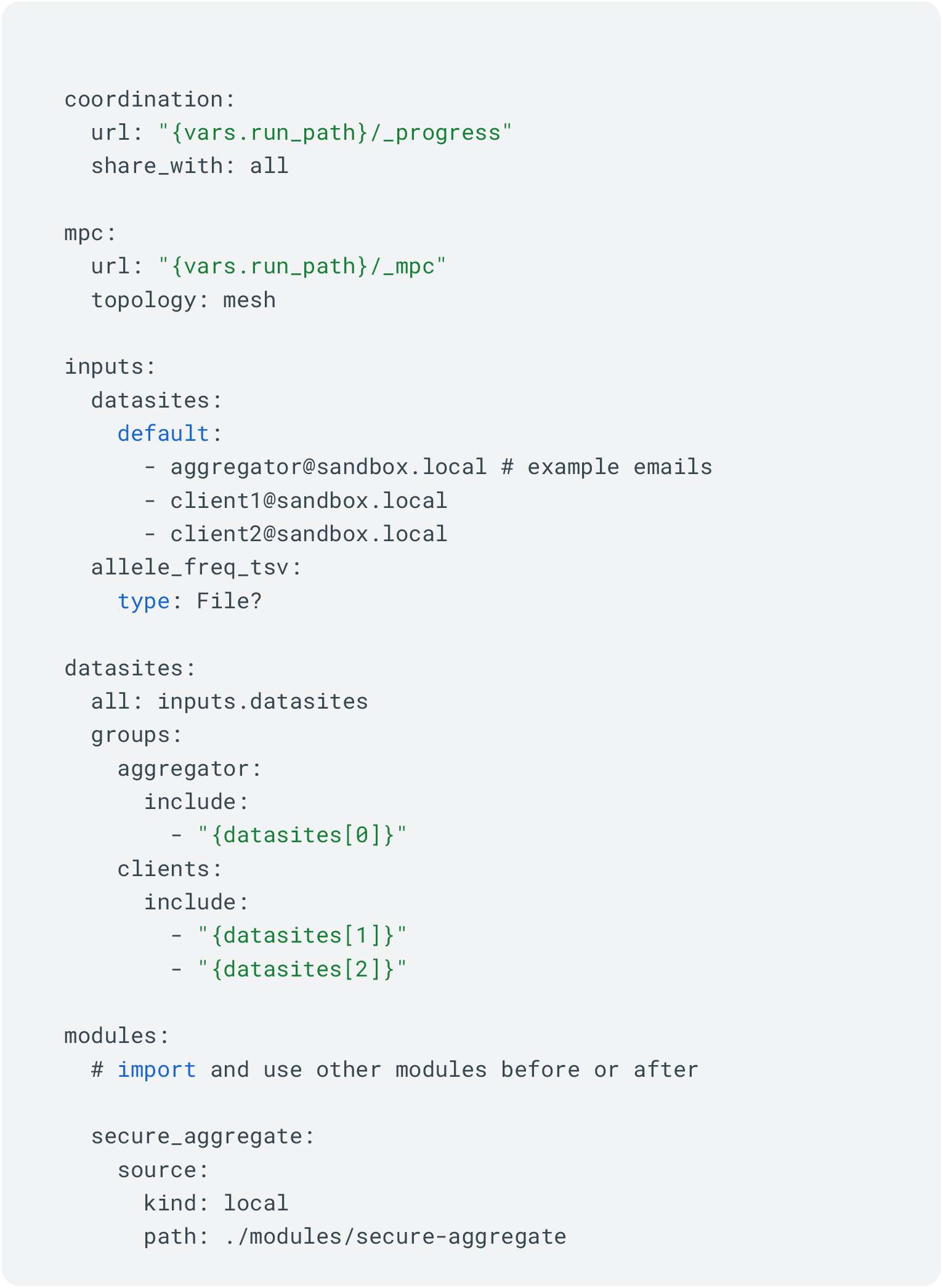
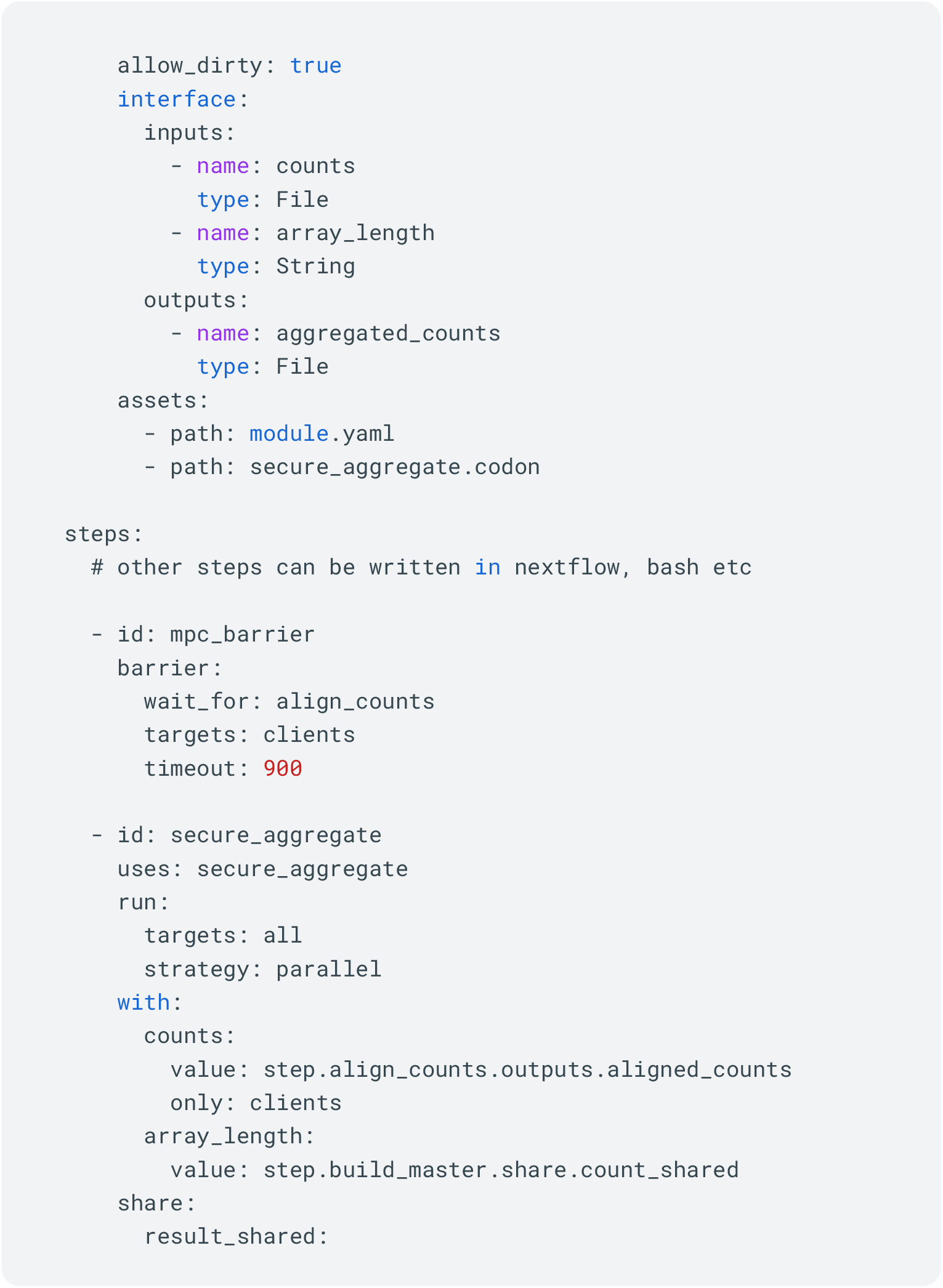
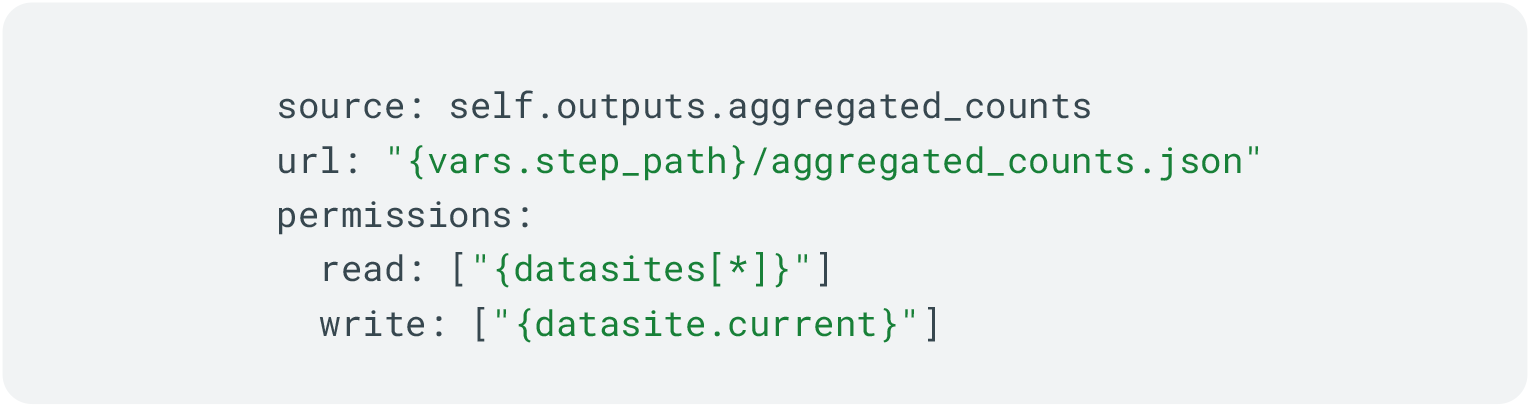

### Syqure SyftBox wrapper for Sequre/Shechi

Sequre natively handles key generation and inter-party communication over standard TCP connections, assuming computing parties are reachable via public IP addresses. In practice, however, BioVault’s target users – including researchers at academic institutions and clinical centers – typically operate behind institutional firewalls or consumer NAT without publicly addressable network endpoints. To enable secure computation between such parties, BioVault wraps Sequre with a zero-configuration proxy that establishes peer-to-peer WebRTC connections between datasites, using Syftbox’s TURN relay server for connection negotiation (Syqure). This allows encrypted inter-party communication to traverse firewalls transparently without requiring users to configure network infrastructure or expose public IP addresses. BioVault further integrates Sequre/Shechi into its pipeline framework (Flows), which allows users to compose multi-step workflows combining standard plaintext computation (via Nextflow or bash) with secure computation steps (via Sequre/Shechi). This design enables researchers to selectively apply cryptographic protection to specific stages of an analysis – for example, performing local preprocessing and quality control in plaintext before executing a secure aggregation step — without rewriting entire pipelines for secure execution. Flows are specified as YAML configurations in which each module declares its execution mode, inputs, and outputs, and BioVault coordinates routing between plaintext and secure steps across datasites. Sequre/Shechi is compatible with MacOS and Linux. To support BioVault’s cross-platform deployment (including Windows), Sequre is executed on Windows within a Linux container managed by the BioVault desktop application, using Docker or Podman as the container runtime. Containerized execution currently operates through BioVault’s file-synchronization layer, which introduces higher latency compared to native TCP or WebRTC; native WebRTC proxy support for containerized Sequre is under development.

### A review of Sequre/Shechi

BioVault integrates secure computation through Sequre^31^/Shechi^32^, an end-to-end, statically compiled framework for building secure multiparty computation (MPC), homomorphic encryption (HE), and multiparty homomorphic encryption (MHE). The framework takes computational pipelines written in Python-like syntax (via the Codon compiler) and compiles them into optimized secure computation programs through static code analysis. During compilation, Sequre/Shechi automatically identifies operations on secure data types, generates a secure expression tree, and applies scheme-specific optimizations — including automatic selection between MPC and HE routines, polynomial restructuring, and prioritization of plaintext over ciphertext operations — to minimize computation and communication overhead without manual optimization by the user.

For MPC, Sequre implements additive secret sharing under an honest-but-curious security model, in which private values are split into cryptographic shares distributed across computing parties such that no individual party can reconstruct another’s inputs. The protocol operates under a server-aided (trusted dealer) model, in which an auxiliary party generates correlated randomness to accelerate computation. For HE and MHE workflows, Shechi (now also called Sequre) extends this foundation with lattice-based homomorphic encryption (based on the CKKS scheme), enabling computation on encrypted data without decryption. MHE combines both approaches by converting between HE ciphertexts and additive secret shares as needed, allowing the compiler to leverage the strengths of each scheme — HE for non-interactive computation and MPC for efficient interactive operations — within a single pipeline. Shechi represents the first MHE compiler and has been shown to achieve up to 15-fold runtime improvements over manually optimized state-of-the-art implementations.

Example.codon SMPC addition

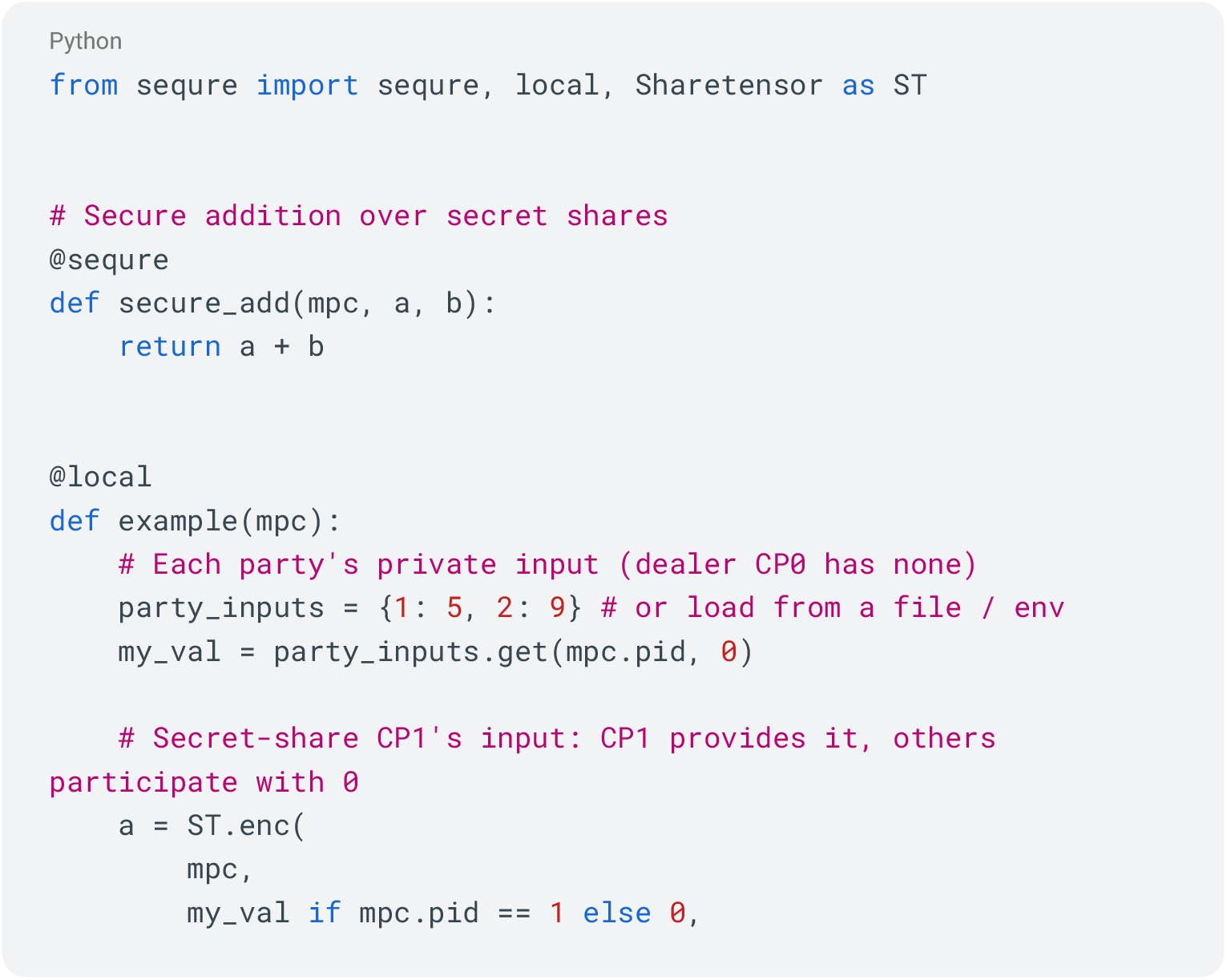
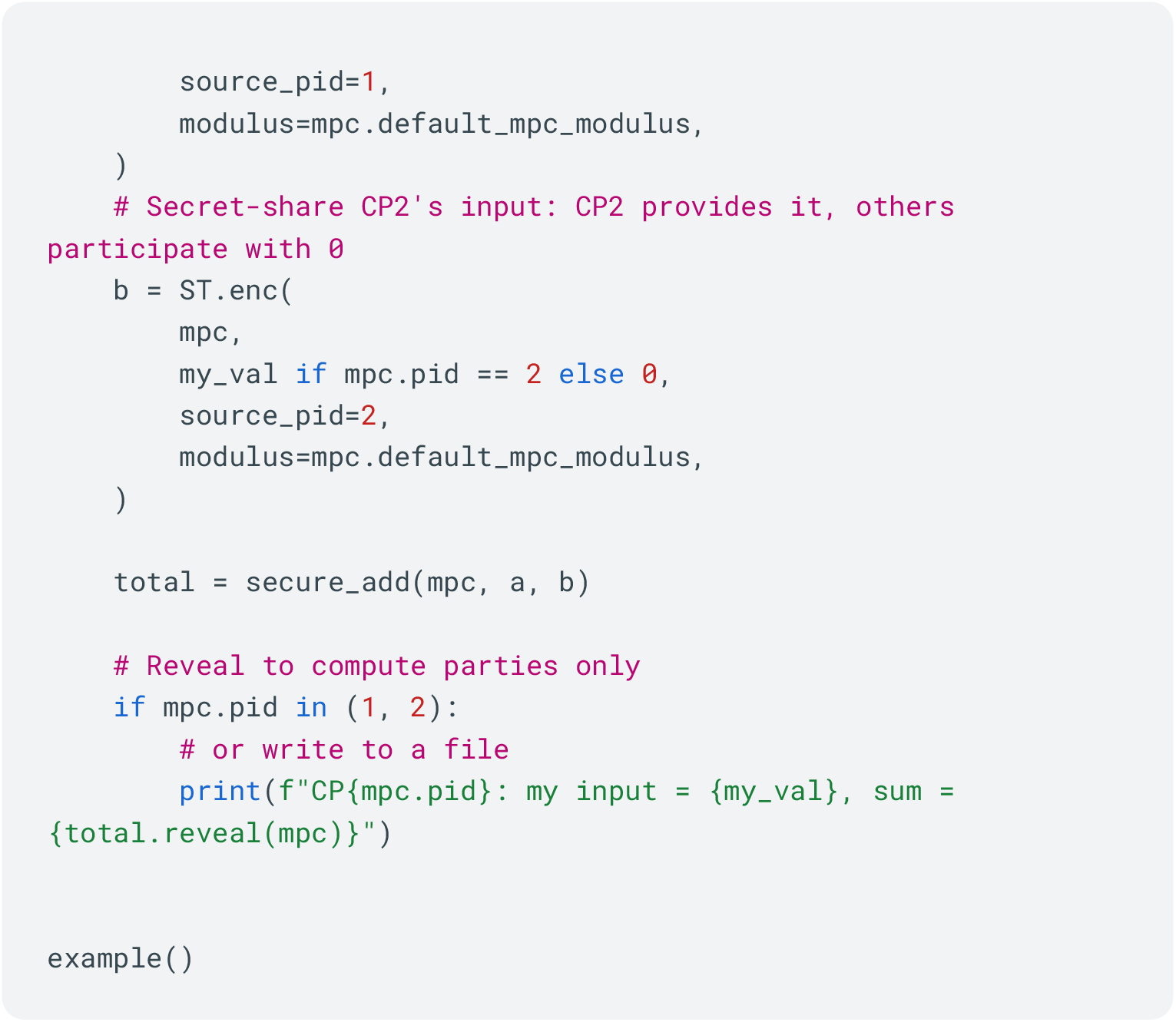

### Usability and Portability

To make this myriad of complex moving parts more user friendly, we built BioVault Desktop a native application for MacOS, Linux and Windows written in Rust and Javascript utilizing the Tauri framework.

The desktop application imports the Biovault CLI shared library code and syftbox-sdk and manages a variety of third party tools such as Nextflow via Java, container runtimes such as Docker or Podman, uv for Python AND Jupyter environment handling and Sequre and Codon via our Syqure wrapper; to provide access to optimized MPC and HE JIT compilation.

The desktop application has a user friendly onboarding process which generates keys for new users showing them a one time BIP39 recovery phrase, handling SyftBox server registration with email 2FA and guiding users through dependency installation with one click installers for common packages.

Installing dependencies and managing PATH on operating systems is complicated even for technical users and can conflict with existing setups.

To make things even easier we bundle several open source binaries with the desktop application which do not pollute users PATH and provide an out of the box experience:

- Java / JRE (open-source)
- Nextflow
- uv for Python
- Syqure (with Sequre and Codon)
- SyftBox client

Users of the desktop application can browse and add other users’ public keys and initialize chat with one or more users.

The chat interface displays some shared objects and results as rich elements with options to save or run code from the chat window. Published datasets can be viewed and subscribed to allowing the user to run pipelines against a local copy of the data for testing.

Data can be loaded via the file open interface and a wizard guides the user through a step by step process designed to optionally detect participant ID information for future referencing.

Common formats currently supported are SNP array.txt files with rsIDs and genotype information (similar to 23andme), VCFs and bam/cram raw files. Users are required to import their own reference genome files and assign them to each file for raw file usage.

Once data is loaded, users can run flows against their data. A flow is a yaml specification which allows for one or more parties to combine inputs in a step by step process of modules resulting in one or more outputs. We provide several simple example flows to test on supported file formats and sample data to use them with.

Each module is a self contained step which can use either Nextflow, Bash or Syqure to execute its instructions. Nextflow itself is configured by default to use containerization to provide portability on different operating systems and cpu architectures. Additionally containerization provides security sandboxing benefits when running untrusted code through restricted file system mounting and network isolation.

We use ‘--network none’ with Nextflow by default so that computations cannot easily exfiltrate data.

Users can optionally choose to publish their data by going through the dataset creation wizard which allows them to create a dictionary of key asset names and assign one or more files to either a single property or a Twin mock / private object. When using Twin’s the public side is published to the server in plaintext for others to download.

To ensure BioVault is future proof we have added a configurable and authenticated local HTTP API which allows other programs or AI agents to control the UI directly while keeping actions visible to a human-in-the-loop user.

Current SOTA AI models and agents are easily able to interact with the API as the method names and inputs exposed via the API schema are described in common API parlance.

However despite these powerful capabilities, we advise caution as prompt injection is currently an unsolved problem and users of BioVault have the Lethal Trifecta: Private Data, Network Access and Untrusted Content (incoming messages or code which can contain prompt injection exploits). See Simon Willison’s famous blog post: https://simonwillison.net/2025/Jun/16/the-lethal-trifecta/.

This is an active area of research and the right use of AI agents for these tools with sufficient sandboxing will bring dramatic improvements to productivity, but for now we suggest caution.

BioVault achieves one of its primary goals of bringing compute to the data by providing compatibility for all popular operating systems and consumer hardware. In the global south the prevalence of Windows is higher than other operating systems and many individuals have affordable low resource machines.

Sadly due to the complexity and cost of developing software for different platforms many standard open source tools are not available on Windows which further drives the centralization of data out of individual desktop systems and into linux environments on the cloud.

Despite this, Biovault and Biovault Desktop are capable of running nearly all features at parity where CPU, RAM and disk space allow. We do this by leveraging Rust for native Windows compatible compilation, and containerization to deliver software like Nextflow and Sequre not currently available for Windows. We leverage DIND (docker in docker) to execute Nextflow in a container and have it launch additional containers as needed. On Windows this can be run with either Docker Desktop via WSL/VM mode or Podman. Despite Podman having some current bugs with file mounts we leveraged it for CI testing where Docker Desktop and WSL are not available in standard CI environments.

Several experiments in this paper were conducted successfully on the Windows version of Biovault Desktop so this is not a theoretical support.

### Experiments

All computational workflows were implemented using Nextflow, Python and a custom cli tool BioSynth designed to efficiently convert SNP array genotype files to VCFs to recover the missing REF values for downstream tasks. Most computations relied heavily on widely used, off-the-shelf bioinformatics libraries and tools. Workflow definitions and source code are publicly available at https://github.com/OpenMined/biovault/tree/main/flows. Jupyter notebooks reproducing the experiments and analytical demonstrations are available at https://github.com/OpenMined/biovault-beaver/tree/main/notebooks.

### Simulation of mock single-cell data

The mock scRNA-seq data were simulated using a Python implementation of Splatter^62,63^. We simulated 36,601 genes across 5,000 cells distributed among 10 clusters. Simulation parameters were configured to ensure clear cluster separation in UMAP visualization. Batch effects and doublets were not included in the simulation.

### Processing and analysis of single-cell data

We analyzed a total of 157,351 freshly collected single cells obtained from breast cancer biopsy samples from 30 patients. The single-cell data was processed using Scanpy^64^. Raw UMI counts were normalized to 10,000 counts per cell using sc.pp.normalize_total, then log-transformed with sc.pp.log1p. The top 5,000 highly variable genes were identified with sc.pp.highly_variable_genes. For visualization, cells were embedded in two dimensions with UMAP (sc.tl.umap) using the top 50 principal components; all other parameters were left at default settings. Cell type annotations were retained from the original study.

### Analysis of ECG heart arrhythmia data

We used the MIT–BIH Arrhythmia Database from PhysioNet, comprising 48 half-hour, two-channel ambulatory ECG recordings from 47 subjects (360 Hz) with cardiologist-curated beat annotations (approximately 110,000 annotated beats in total). For heartbeat classification, annotation labels were grouped into five categories: normal (N), fusion (F), supraventricular ectopic beat (SVEB), ventricular ectopic beat (VEB) and unknown (Q). ECG waveforms were converted into beat-level, fixed-length examples using the reference annotations, and a feature matrix (X) with corresponding labels (y) was generated within the data-owning environment and provided as input to downstream classifiers.

For XGBoost^65^ classifier training, we split data into train and test sets using sklearn.model_selection.train_test_split (test_size=0.2, random_state=42). To preserve class balance we used stratified splitting (stratify=y) when all classes contained at least two samples (otherwise stratify=None). We trained an xgboost.XGBClassifier (n_estimators=200, learning_rate=0.05, max_depth=6, scale_pos_weight=1, random_state=42, eval_metric=“mlogloss”). All other XGBoost parameters were left at library defaults. The model was fit once on the training set (model.fit(X_train, y_train)), and test accuracy was computed as the fraction of correct predictions on the held-out test set.

For confusion-matrix evaluation, we predicted test labels with model.predict(X_test) and computed a confusion matrix using sklearn.metrics.confusion_matrix, restricting axes to labels present in either the true or predicted test labels. Confusion matrices were visualized with sklearn.metrics.ConfusionMatrixDisplay.

For t-SNE visualization, we randomly subsampled up to 3,000 examples for speed using a fixed RNG seed (np.random.default_rng(42)), standardized features with sklearn.preprocessing.StandardScaler, and embedded the standardized feature vectors using sklearn.manifold.TSNE (n_components=2, perplexity=30, learning_rate=200, random_state=42, max_iter=1000).

For the Keras MLP baseline, we standardized inputs with sklearn.preprocessing.StandardScaler and one-hot encoded labels with keras.utils.to_categorical. We again used sklearn.model_selection.train_test_split with test_size=0.2, random_state=42, and stratification conditional on minimum class size as above. The Keras Sequential network consisted of a 128-unit Dense layer with ReLU activation, a dropout layer (rate 0.3), a 64-unit Dense layer with ReLU activation, a second dropout layer (rate 0.3), and a final Dense layer with num_classes units and softmax activation. Models were compiled with Adam (optimizer=“adam”) and categorical cross-entropy (loss=“categorical_crossentropy”) and trained for epochs=10 with batch_size=64, validation_split=0.2, and verbose=2. Test accuracy was obtained via model.evaluate(X_test, y_test) and a confusion matrix was computed from argmax class predictions.

For remote execution, both training routines were wrapped as Beaver-executable functions and executed under a data-visitation workflow: the training code was developed and validated on mock data, then submitted for private execution in the data-owning environment; only aggregate outputs (accuracy, confusion matrix, and serialized model weights) were returned to the data scientist. Model parameters training was adopted from Kaggle (https://www.kaggle.com/code/srijitsengupta06/ecg-arrhythmia-classification-mitbih/notebook).

### Segmentation of Brain MRI data using deep learning

We used the Multimodal Brain Tumor Segmentation Challenge 2018 (BraTS 2018) dataset^39,66,67^. BraTS 2018 provides multimodal MRI scans (T1, T1c, T2 and FLAIR). Tumor subregions were manually annotated, comprising GD-enhancing tumor (ET; label 4), peritumoral edema (ED; label 2), and necrotic and non-enhancing tumor core (NCR/NET; label 1). We applied a pretrained BraTS model distributed as the BraTS MRI segmentation bundle from MONAI^68,69^ (https://project-monai.github.io/model-zoo.html#/model/brats_mri_segmentation). The bundle is trained on BraTS 2018 to segment three nested subregions (ET, TC, WT) from four aligned MRI modalities (T1c, T1, T2, FLAIR).

Segmentation inference was performed using a Nextflow pipeline that processes cases independently and executes a fixed sequence of steps: (i) gather the four MRI modalities (T1, T1c, T2, FLAIR) for each case and provide them as inputs to the inference step, (ii) run the pretrained MONAI BraTS bundle in inference mode using the bundled preprocessing and sliding-window configuration (roi_size=[240,240,160], overlap=0.5, sw_batch_size=1, AMP enabled), and (iii) apply post-processing to generate discrete segmentations by thresholding at 0.5 and writing compressed NIfTI masks (.nii.gz, uint8). The pipeline outputs per-case predicted segmentation volumes and optional derivative artifacts, including 2D slice overlays for qualitative inspection and per-case summary tables, enabling batch application of a fixed pretrained model to new datasets without modifying model weights.

### Analysis of rare disease whole genome sequencing data

Whole-genome sequencing (WGS) data were obtained from Sequencing.com. Variant calling and annotation were performed using the Nextflow Sarek^70^ pipeline (nf-core/sarek), a reproducible workflow for WGS data analysis. The pipeline was used for read alignment, quality control, and small variant (SNV/indel) calling, followed by functional annotation. Copy number variant (CNV) analysis was performed separately using a custom Python script developed in-house to detect and prioritize candidate CNVs from WGS data.

### GWAS on Type II Diabetes in Circassian and Chechan Populations

The genotype dataset comprised a merged cohort of 284 individuals in total, distributed across two populations: Circassian (n=139 samples) and Chechen (n=145). Genotyping was performed using the Illumina Infinium OMNI-Express BeadChip platform at the Center for Applied Genomics at the Children’s Hospital of Philadelphia. We received quality-controlled data that had undergone sample-level filtering (call rate >95%, heterozygosity within five standard deviations, unrelated individuals) and SNP-level filtering (call rate >95%, minor allele frequency >1%, Hardy-Weinberg equilibrium p-value >10^−4^ in controls). The merged dataset comprised 639,942 autosomal SNPs. For the GWAS analysis, we selected Type 2 Diabetes (T2D) as the target phenotype, with 67 cases and 215 controls (2 phenotypes missing). To account for population stratification, we first generated a subset of 115,852 independent SNPs, via LD pruning using a pairwise approach (window size 50 SNPs, step 5 SNPs, r^2^ threshold 0.2). We further excluded 330 ambiguous AT/GC SNPs, resulting in 115,522 SNPs for principal component analysis. We computed the top 10 principal components and included them as covariates in logistic regression-based association testing across all 639,942 quality-controlled SNPs. All steps were carried out using PLINK^43^.

### Calculation of allele frequencies across Caribbean populations

To enable cross-border population genetic analysis while preserving data sovereignty, we performed remote allele frequency calculations using the BioVault decentralized framework. This “data visitation” workflow allowed computations to be initiated from a central site (United States) and executed locally at remote data sites in Saint Lucia and Bermuda. Per-country SNP-array callsets—encompassing cohorts from the Bahamas (n = 168), Barbados (n = 125), the British Virgin Islands (n = 106), Bermuda (n = 488), Saint Lucia (n = 97), and Trinidad and Tobago (n = 76)—were processed locally into individual VCF files. These VCFs were then processed into normalized long-row format to obtain cohort-level allele counts across all variants. These aggregate statistics (allele counts, total alleles, and calculated frequency) were returned to the initiator, ensuring that no individual-level genotype data was transferred across borders. This entire process– from VCF generation to computing aggregate– statistics was done using a containerized Nextflow (DSL2) workflow (BioVault allele-freq v0.1.0) built on the Rust-based BioSynth (bvs) framework using allele reference data from NCBI dbSNP.

For comparative benchmarking, we downloaded reference data from gnomAD (v4.1) and extracted allele frequencies for global and ancestry-specific populations, including African (AFR), non-Finnish European (NFE), and South Asian (SAS) panels. Relationships between Caribbean and reference populations were assessed using Pearson correlation coefficients (r) and visualized via two-dimensional binned scatter plots. Targeted analysis was further performed on a curated panel of pharmacogenetically and clinically relevant variants to identify population-specific enrichment or depletion relative to gnomAD reference frequencies.

### Population-level allele frequency calculation, via secure multiparty computation

To demonstrate secure federated computation compatibility within the BioVault network, we performed secure aggregation of allele frequencies across datasites in the Caribbean (Bermuda and Saint Lucia) using MPC. The protocol was executed across three BioVault desktop instances: two datasites holding private genotype data (corresponding to the Caribbean cohorts described in **Figure 4**) and one aggregator (remote server) acting as the trusted dealer **(Figure 5A)**.

Each datasite independently generated a locus index containing variant identifiers from its local callset. These indices were shared with each datasite to build a shared variant index. A subset of 100K SNPs from the shared subset were chosen for allele frequency estimation. Each datasite then locally aligned its per-variant allele count and total allele number vectors to this subset, padding any missing variants with zeros. Per-site allele count and total allele number vectors were turned into encrypted shares (secret-sharing based MPC). These shares were securely aggregated to produce total allele counts and numbers, and then securely divided. The final result is an encrypted share of allele frequency estimation. At no point during the secure computation did any party gain access to another site’s per-variant allele counts, sample sizes, or allele frequencies; only the final pooled totals were revealed upon share reconstruction.

The computation was performed over the live BioVault network with inter-site latencies exceeding 300 ms. Secure aggregation of 200K data points (allele count and allele number values across the shared variant set) completed in approximately 20 minutes (Figure 5C), representing roughly 4-fold overhead compared to localhost TCP benchmarking. Concordance between securely aggregated and non-secure centralized allele frequencies was assessed by direct comparison (Figure 5B).

## Acknowledgements

We thank members of the OpenMined Foundation for helpful discussions. We thank F. Crawley, S. Edmunds, M. Jay, G. Church, Y. Li, W. Liu, T. Pollard, J. Reichardt, and R. Taft for insightful discussions. We thank M. Jay for enabling the use of his genome data for this study. Funding was provided in part by the OpenMined Foundation.

## Author contributions

X.D.C., M.J., R.D., and C.W. conceived the study. M.E., T.P.T., M.J., Z.A., D.F., R.D., C.W., and X.D.C. designed and performed the experiments and analyzed the data. M.J. and K.J. developed the software platforms. M.E., T.P.T., M.J., K.J., and X.D.C. wrote the manuscript with input from all authors.

## Declaration of Interests

M.E. and M.J. are employed by, or have received compensation from, the non-profit OpenMined Foundation. All other authors declare no competing interests.

### Declaration of generative AI and AI-assisted technologies in the writing process

During the preparation of this work, the author(s) used ChatGPT in order to improve readability and language clarity. After using this tool or service, the authors reviewed and edited the content as needed and take full responsibility for the content of the publication. During the production of code for BioVault and its libraries, we utilized SOTA AI coding models and harnesses from Anthropic and OpenAI.

### Ethics statement

All human genomic data used in this study, including the Jordan and Bermuda datasets, were collected under protocols approved by the relevant institutional review boards (IRBs) in their respective jurisdictions, with informed consent obtained as required. Analyses were conducted using a data visitation framework in which private data remained within the secure environment of the data owners. Computational requests were executed locally by the data owners, and only results that were reviewed and determined not to contain private or identifying information were approved for release. No raw or private data were transferred to external investigators.

This study also includes genomic data from a single individual who is a co-author of this work. Explicit informed consent was obtained from this individual for analysis and publication under the same data visitation framework described above.

Because peer-to-peer data visitation models involve remote execution of analytical code without direct data transfer, their classification under existing IRB and data governance frameworks is not always explicitly defined. We consulted with institutional advisors and domain experts regarding the ethical and regulatory implications of this approach. All analyses were conducted in accordance with applicable local regulations and data use agreements. We recognize that computational architectures for data visitation represent an evolving policy area and welcome continued discussion to clarify regulatory guidance and best practices.

## Data availability

The scRNA-seq data of metastatic breast cancer biopsies is publicly available from the Single-Cell Portal (https://singlecell.broadinstitute.org/single_cell/study/SCP2702) and interactively browsable through CELLxGENE (https://cellxgene.cziscience.com/collections/a96133de-e951-4e2d-ace6-59db8b3bfb1d). The MIT-BIH Arrhythmia Database is available from Physionet (https://physionet.org/content/mitdb/1.0.0/). The BraTS 2018 dataset is available on kaggle (https://www.kaggle.com/datasets/sanglequang/brats2018/data).

## Code availability

BioVault is available as a free, open-source desktop application at https://github.com/OpenMined/biovault-desktop. This uses the core BioVault CLI (https://github.com/OpenMined/biovault) via Tauri, a framework for building cross-platform applications (https://github.com/tauri-apps/tauri).

The underlying networking layer is powered by SyftBox (https://github.com/OpenMined/syftbox; https://www.syftbox.net), a protocol for building and federating privacy-preserving computations across distributed networks, with its corresponding SDK providing the End-to-end encrypted communication in BioVault (https://github.com/OpenMined/syftbox-sdk). The SyftBox environment manager, sbenv (https://github.com/OpenMined/sbenv), enables isolated multi-site testing. Reference data files are available at https://github.com/OpenMined/biovault-data

Secure multi-party computation for privacy-preserving aggregation across sites is performed using Syqure (https://github.com/madhavajay/syqure), a Rust wrapper around the Codon/Sequre MPC compiler (https://github.com/0xTCG/sequre).

BioVault Beaver (https://github.com/OpenMined/biovault-beaver) is the Python bioinformatics toolkit which provides the allele frequency and genomic analysis pipelines used in this study. Tutorials to perform twin analyses from data scientist or data owner perspectives are available in the Beaver repository (https://github.com/OpenMined/biovault-beaver/tree/main/notebooks).

The MRI brain tumor segmentation model is available as a MONAI bundle (https://project-monai.github.io/model-zoo.html#/model/brats_mri_segmentation).

BioScript (https://github.com/OpenMined/bioscript) offers additional multi-language bioinformatics utilities. Synthetic genotype data for testing and development was created using BioSynth (https://github.com/OpenMined/biosynth).

