## Supplementary Information for "BioVault: A privacy-first data visitation platform for equitable global collaboration in biomedicine"

**Supplemental Information**

Supplementary Note 1: Description of Syftbox
Supplementary Note 2: Description of BioVault Beaver
Supplementary Note 3: Description of CariGenetics Datasets

Supplementary Note 1: Description of Syftbox

SyftBox (https://www.syftbox.net/) is an open-source protocol developed by OpenMined Foundation (https://openmined.org/) that enables secure, decentralized coordination of remote computation across independently governed environments. Participants are organized as datasites– local directories on individual machines that act as nodes in a distributed file and compute network. Each datasite is fully owned and operated by its host, and data never leaves the local environment unless explicitly authorized.

The protocol follows a three-layer architecture:

- Application layer- Supports applications written in any programming language via a custom syft:// URL scheme.
- Communication layer – Provides event-driven remote procedure calls (RPC), HTTP-to-filesystem bridging, object serialization, and end-to-end encryption.
- Infrastructure layer – Includes an optional relay cache server that forwards encrypted data between peers without requiring direct TCP connections.

Identity is managed through email-based two-factor authentication that binds users to public-private key pairs conforming to W3C Decentralized Identifier (DID) standards. This enables mutual authentication and asynchronous encrypted communication between participants.

End-to-end encryption is optionally implemented using a simplified Extended Triple Diffie-Hellman (X3DH) protocol. The implementation reduces the exchange to two Diffie-Hellman operations while preserving forward secrecy through ephemeral keys and mutual authentication via signed prekeys.

File synchronization between datasites is mediated by a public relay cache server that stores and forwards encrypted payloads, enabling asynchronous collaboration without requiring simultaneous connectivity. The relay server operates on open-source code, and the protocol supports independent operation of additional relay servers.

Access control follows a Unix-style permission model with read, write, create, and admin permissions specified per file path and per user identity in declarative YAML configuration files. Datasite owners retain irrevocable administrative authority over their data.

SyftBox is transport-agnostic. WebSocket connections support low-latency communication for interactive protocols such as secure multi-party computation (MPC), while HTTP polling supports longer-timescale synchronization. Data can also be transferred via USB (sneakernet), FTP, or cloud-drive synchronization, enabling deployment in air-gapped or high-security environments.

Computation can follow either a data visitation model, in which analysis code executes locally within the data owner’s environment and only approved results are returned, or a federated protocol where data exchange occurs across defined topologies and steps.

SyftBox is implemented in Go, with Python and JavaScript SDKs, and is released under the Apache 2.0 license (current release: v0.8.7).

Supplementary Note 2: Description of BioVault Beaver

BioVault Beaver (https://github.com/OpenMined/biovault-beaver) is a Python framework designed to enable collaborative analysis of sensitive genomic and biomedical data without transferring raw private datasets. The system implements a data visitation model, in which analytical logic moves to the data owner's environment, while private data remains local.

Importantly, unlike many static upfront remote workflows, Beaver is designed for iterative eager execution to facilitate the crucial exploratory Quality Control and ETL processes required during the preliminary stages of data science.

**Twins: A Dual-Value Data Abstraction**

The core abstraction in Beaver is the Twin, a dual-value Either Type containing:

- A public (mock) value, used by collaborators for method development and exploratory analysis.
- A private (real) value, kept private within the data owner's environment, but transferable upon approved release to the data scientist

Collaborators develop workflows against public representations that preserve structure and schema. When analysis is finalized, computation is executed on the private value within the owner's environment. This approach separates algorithm sharing from data sharing.

**Structured Serialization and Controlled Loading**

All shared objects are serialized into structured .beaver envelopes containing metadata and version information. Serialization explicitly disallows unsafe pickle-based execution.

For large datasets, Beaver references external artifacts rather than embedding raw data directly. File loading is mediated by controlled loader descriptors that restrict imports and validate file paths, reducing risk during deserialization and execution.

Registry-Based Discovery

Beaver provides a per-user registry that catalogs shared public objects. Public representations can be discovered and lazily loaded by collaborators without exposing private data. Private components are never published to the registry.

This mechanism enables reproducible discovery of datasets and variables while maintaining strict separation between public metadata and private content.

**Encrypted Storage and Sessions**

When integrated with encrypted storage backends (e.g., SyftBox), shared envelopes and artifacts are encrypted for intended recipients. Cross-user data exchange occurs through explicit session workflows, in which:

1. A requester submits a computation request.

2. The data owner executes the computation locally on private data.
3. The owner reviews and approves outputs before release.

Only approved results are returned. Private data never leave the owner's environment.

**Remote Computation Model**

Beaver supports structured computation requests in which analytical functions and input references are packaged and transmitted to the data owner. Execution occurs locally, and results are returned as controlled outputs. Public mock outputs on the requester side remain unchanged, preserving reproducibility.

This model allows collaborators to validate code logic without direct access to sensitive data, enabling privacy-first statistical analysis, visualization, and model evaluation.

**Security Model**

Security is enforced at multiple layers:

- No direct TCP connection between parties
- Controlled deserialization with module/type restrictions
- Blocking of arbitrary function execution by default
- Path sanitization and URL validation
- Optional restricted execution environments
- Encryption of cross-user artifacts

Together, these mechanisms implement a layered security architecture appropriate for regulated genomic and biomedical data.

**Conceptual Summary**

BioVault Beaver combines dual-value data objects (twins), structured serialization, encrypted transfer, and explicit approval workflows to enable privacy-first eager execution on remote private data. This framework integrates with standard tools such as Jupyter Notebooks and remains fully compatible with the Python data science ecosystem. By separating public analytical development from private data execution, Beaver operationalizes a controlled data-visitation framework suitable for sensitive genomic datasets, multi-institutional studies, and federated biomedical research.

Supplementary Note 3: Description of CariGenetics Datasets

**Research Cohort**

The Caribbean Prostate Cancer Genetic Study is a cross-sectional study of patients with prostate cancer and non-patients born in the Caribbean. Recruitment into the study was conducted from November 2024 to December 2025, with data analyzed between June 2025 and January 2026. A total of 248 participants distributed across two populations, Bermuda (n=158 samples) and Saint Lucia (n=90 samples), were enrolled. All these individuals provided explicit written consent, allowing their de-identified individual-level research information to be used outside of CariGenetics in research collaborations. Ethics approval was obtained by the BHB Research Ethics Committee in Bermuda and the Medical & Dental Council (Saint Lucia W.I.) Research Ethics Committee in Saint Lucia. The study was led by a principal investigator (C.W.). Participants were identified by treating physicians and media outlets, including radio, newspaper, and television advertisements. Individuals needed to have all 4 grandparents born in the Caribbean; the ability to provide consent, and the ability to provide a blood specimen. All participants gave informed consent, which included presentation of results in publication.

Following written informed consent, an EDTA whole blood sample was obtained. Genomic DNA was isolated following the manufacturer's instructions for the NEB Monarch HMW Blood & Cells Kit in Bermuda. Aliquoted DNA samples were shipped to Dynamic DNA Laboratories in MO, USA, for genotyping.

**Commercial Cohort**

Commercial direct-to-consumer testing was conducted from May 2024 to October 2025. A total of 931 people distributed across six populations, Bahamas (n=168 samples), Barbados (n=126 samples), Bermuda (n=448 samples), BVI (n=106 samples), Saint Lucia (n=7 samples), and Trinidad and Tobago (n=76 samples) gave optional explicit written consent providing permission for their anonymized data to be used by CariGenetics for internal and external research. Only summary statistics were shared externally; no individual-level data were released outside CariGenetics. Following written informed consent, a cheek swab sample was obtained and shipped to Dynamic DNA Laboratories in MO, USA for genotyping.

Genotyping was performed using the Illumina Infinium Global Screening Array v3 with DTC Booster content. Genotype calling was performed using a laboratory-validated custom cluster file (.egt) generated in accordance with Illumina-recommended best practices and optimized using high-quality reference samples. Sample-level quality control was conducted before data release and included filtering for overall genotype call rate (>95%), concordance between reported and genetically inferred sex, and probe intensity metrics, with samples required to have a p10GC value ≥0.40. Any additional variant- or cohort-level filtering was performed downstream by the investigators.
